# Coding triplets in the tRNA acceptor-TΨC arm and their role in present and past tRNA recognition

**DOI:** 10.1101/559948

**Authors:** Ilana Agmon, Itay Fayerverker, Tal Mor

**Affiliations:** Institute for Advanced Studies in Theoretical Chemistry, Schulich Faculty of Chemistry - Technion - Israel Institute of Technology, Haifa, Israel; Fritz Haber Research Center for Molecular Dynamics, Hebrew University Jerusalem, Israel; Department of Computer Science, Technion, Haifa, Israel

**Author notes:** Corresponding author: Ilana Agmon, Institute for Advanced Studies in Theoretical Chemistry, Schulich Faculty of Chemistry - Technion - Israel Institute of Technology, Haifa 3200003, Israel.

**Keywords:** Aminoacylation, Genetic code, Origin of life, Synthetase, tRNA, Translation

## Abstract

The mechanism and evolution of the recognition scheme between key components of the translation system, i.e., tRNAs, synthetases and elongation factors, are fundamental issues in understanding the translation of genetic information into proteins. Statistical analysis of bacterial tRNA sequences reveals that for six amino acids, i.e. for Ala, Asp, Gly, His, Pro and Ser, a string of 10 nucleotides preceding the tRNA 3’end, carries cognate coding triplets to nearly full extent. The triplets conserved in positions 63-67 are implicated in the recognition by EF-Tu, and those conserved in positions 68-72, in the identification of cognate tRNAs and their derived minihelices, by class IIa synthetases. These coding triplets are suggested to have primordial origin, being engaged in aminoacylation of prebiotic tRNAs and in the establishment of the canonical codon set.

## Introduction

tRNA is a vital component of the translation system due to its role in linking the genetic code with the synthesized protein. The modern tRNA is an L-shaped molecule. One end of the molecule is the universally conserved single stranded NCCA tail, which carries the cognate amino acid (aa), while at the other end, about 75Å away, the anticodon (AC) loop that fully characterizes the attached amino acid is located.

The mature tRNA interacts with three key components of the translation system - the cognate aminoacyl-tRNA synthetase (aaRS) that specifically aminoacylates it; the elongation factor (EF-Tu in bacteria) that accommodates the aa-tRNAs and carries them, in a ternary complex, to the ribosome; and the ribosome where the attached amino acid is incorporated into the growing polypeptide. The low overall frequency of amino acid misincorporation in translation (10^−3^ to 10^−4^) reflects the cumulative fidelity of these three principal reactions [1].

Recognition of aa-tRNA on the ribosome is accomplished via the interactions of the mRNA codon accommodated on the small ribosomal subunit, with the AC loop. EF-Tu, which carries a variety of different tRNAs, recognizes each of them specifically. The minimal fragment of tRNA that can interact efficiently with EF-Tu consists of a 10 base-pair helix from the acceptor-TΨC stem, linked to the aa-3’end tail [2], from which nucleotides 63–67 form the contact interface [3].

Aminoacyl-tRNA synthetases are divided into two distinct classes; Class I synthetases possess active sites with the Rossmann fold motif, while class II have active sites composed of antiparallel β-sheets with three highly conserved motifs; 1,2 and 3 [4]. Usually there is one synthetase for each amino acid, 10 of them belonging to class I and 10 to class II, but LysRS has a class II form in addition to the canonical class I synthetase [5]. Synthetases commonly acquire the identity of their cognate tRNAs from the anticodon stem-loop. Yet, this loop is not always the principal determinant for aminoacylation [6,7] and in some cases no physical contact is made between the aaRS and the anticodon loop [8]. In several tRNAs, major identity elements are found in the core of the molecule, as is the case, for example, with the large variable loop in tRNA^Ser^. An additional set of identity elements classified as ‘‘determinants’’ and ‘‘anti-determinants’’ [9], such as the discriminator base (N73) [10] and specific nucleotides in the acceptor stem [11-13], were shown to participate in the tRNA recognition by the synthetase.

The recognition determinants embedded in the first base pairs of the tRNA acceptor stems, as well as in RNA minihelices derived from their cognate acceptor-TΨC stems (also known as “tRNA acceptor branches”), were demonstrated to carry coding information sufficient for their specific aminoacylation [9,11-16]. This non-AC aminoacylation process was suggested to be controlled by an “operational code” [12] that relates few nucleotides in the acceptor stem to specific amino acids. Its mode of action was proposed to be a remnant of an early mechanism for specifically aminoacylating proto-tRNAs by proto-synthetases, possibly associated with a second code [12,17], distinct from the nucleotide triplets of the standard genetic code [16].

Here we analyze large sequence data sets from bacterial tRNA genes and report the occurrence, to a very high extent, of cognate coding triplets in specific locations of the acceptor-TΨC arm belonging to six amino acids; Ala, Asp, Gly, His, Pro and Ser. Coding triplets conserved in positions 63-67 are implicated in the tRNA recognition by EF-Tu, while the triplets conserved in the nucleotide range 68-72, are proposed to be involved in the contemporary tRNA recognition by synthetases of class IIa. In the evolutionary context, these conserved coding triplets are suggested to have served as identifiers of proto-tRNAs in the primordial aminoacylation process, and to have had a primal role in the emergence of the contemporary codon set.

## Results

In bacterial tRNAs, the sequence of the string preceding the 3’ end (pre-3’end), ranging from nucleotide 63 to 72 is diverse [18], even among different strains of the same bacterium (Table 1, File S1). Statistical analysis was performed on large scale sequence data of the pre-3’end strings, acquired from the tRNAdb database [19,20], http://trna.bioinf.uni-leipzig.de, which holds 128 bacterial tRNA sequences obtained from elongator-tRNAs involved in protein synthesis, and over 6200 bacterial tRNA genes, belonging to the 20 canonical amino acids. To avoid the inclusion of pseudo-tRNA genes, i.e., those not coding for elongator-tRNAs, only tRNA gene-sequences having a full 3’ end and a consensus discriminator base [10] were incorporated into the statistics (Materials and Methods).

**Table 1.**
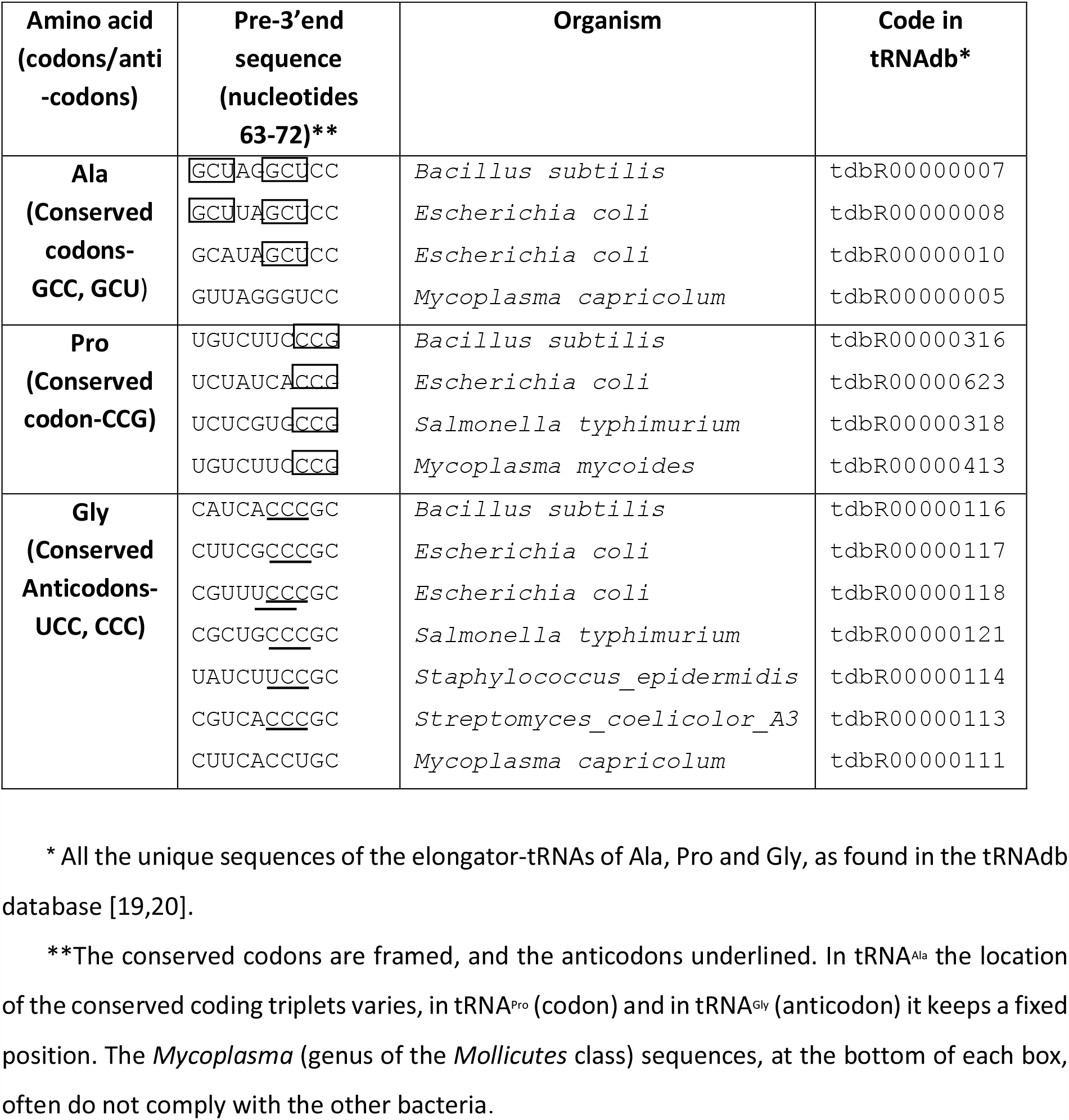
Conserved stem-codons and stem-anticodons in the tRNA Pre-3’end sequences

When read from the 5’ to the 3’, these strings, in the tRNAs of nine amino acids, were found to hold excessive cognate codon and/or anticodon triplets, termed here sC/sACs, i.e. stem-codons and/or stem-anticodons (Tables 1, 2, Table S1a, File S1). One class of bacteria, the *Mollicutes*, deviated from the statistical behavior exhibited by the other bacteria (Table 1, File S1, Table S2), and its exclusion resulted in a sample of 3324 tRNA gene-sequences from over 100 bacterial species. Statistical analysis that incorporated the 350 tRNA-genes of *Mollicutes* did not differ significantly from the results obtained with the data sample used in the current study.

**Table 2.** Statistics of the conserved sC/sACs in the pre-3’end strings from bacterial tRNA genes

| Amino acid*<br>(no. sequences) | Conserved sC/sAC set* | | Observed Occurrence** | Expected occurrence***<br>($\pm 0.8\%$) | *P* |
| --- | --- | --- | --- | --- | --- |
|  | Codons | Anticodons |  |  |  |
| **Ala** (209) | **GCU, GCC** |  | 83 % | 50 % | E-22 |
| **Asp** (110) |  | **GUC** | 96 % | 17 % | E-106 |
| **Gly** (243) |  | **CCC, UCC** | 94 % | 57 % | E-31 |
| **His** (83) | **CAC, CAU** |  | 90 % | 22 % | E-51 |
| **Pro** (180) | **CCG** |  | 98 % | 37 % | E-64 |
| **Ser** (290) | **UCA, UCC, UCU** |  | 99 % | 46 % | E-72 |
| Arg (305) | **CGG, AGG**<br>CGA, CGC, CGU | CCU | 95 % | 84 % | E-7 |
| Leu (366) | CUN | CAG | 84% | 63% | E-17 |
| Val (201) | GUN | NAC | 88 % | 58 % | E-17 |

\* Cognate triplets whose conservation is considered as robust, as well as the robustly conserving amino acids (Materials and Methods) are depicted by bold letters.
\*\* % of pre-3' end strings holding at least one sC/sAC from their conserved set at any location along the string.
\*\*\* Derived from random oligonucleotides simulation with "sample" distribution (A,C,G,U)=(10%,46%,24%,20%). The statistics for the other two distributions and the statistics for the distributions calculated for the remaining 11 amino acids are given in Tables S1a and S1b, respectively.
N- any nucleotide. *P*- the probability that the observed occurrence of a particular sC/sAC set is random. $P \leq 0.01$ is considered statistically significant.

### A group of nine conserving amino acids

The occurrence of the cognate codons and anticodons in the pre-3’end strings of the 20 canonical amino acids was determined. Two distinct groups with a wide gap between their cognate sC/sAC occurrences were observed: the pre-3’end strings of the members of the first group, that is, of Ala, Arg, Asp, Gly, His, Leu, Pro, Ser and Val, were found to contain cognate sC/sACs in over 85% of the strings of each amino acid. These amino acids were denoted “conserving amino acids”. For the remaining 11 “non-conserving amino acids”, i.e., for Asn, Cys, Gln, Glu, Ile, Lys, Met, Phe, Thr, Trp, Tyr, only 0%-40% of their pre-3’end strings carried a cognate sC/sAC (Table S1b).

For each of the amino acids of the first group, the set of specific coding triplets that accounts for 98% of the sC/sAC occurrence within the strings carrying any cognate sC/sAC, is denoted the “conserved sC/sAC set”, and the codons and anticodons belonging to any conserved set are named “conserved sC/sACs” (Table 2) (Material and Methods). These conserved sC/sACs are present in 83% or more of the pre-3’end strings of each conserving amino acid and they typically constitute only a subset of the corresponding standard codons and anticodons table (Table 2). Interestingly, the cognate sC/sACs conserved in the pre-3’end strings are often not correlated with the anticodon triplet in the AC loop of the same tRNA (File S1).

For each of the nine conserving amino acid, the observed occurrence of conserved coding triplets in their pre-3’end was compared with its expected occurrence. The expected occurrences were obtained by analyzing a large dataset of random 10-mer oligonucleotides. The construction of the randomized sequences and the estimation of the expected occurrence of a specific set of sC/sACs is described in the Materials and Methods section. The randomized strings were constructed with three nucleotide distributions: 1. Distribution identical to the average found in all the bacterial pre-3’end strings analyzed (“sample” distribution), i.e.: A=10%, C=46%, G=24%, T=20%; 2. Equal distribution of the four nucleotides; 3. The individual nucleotide distribution found in the pre-3’end string of each amino acid (Tables S1a, b).

Analysis of the observed vs. expected sC/sAC occurrences in the data of the conserving amino acids confirmed that the observations are clearly nonrandom, i.e. that the conserved sC/sACs of the nine conserving amino acids show observed occurrences that go far beyond the statistical expectation (*P*<<0.01), a result obtained with all three distributions (Table 2, Table S1a). For the non-conserving amino acids, in the absence of a conserved sC/sAC set, only the combined occurrence of all the cognate sC/sACs was checked. The combined occurrences were found to lie in the range 0-40% for each amino acid, which, for Gln, Glu, Lys, Met, Thr and Tyr is less than statistically expected. For the remaining five amino acids, it practically accords with the statistical expectations (Table S1b). Noteworthy, the coding triplets in the Thr pre-3’end strings demonstrate idiosyncratic features - it has the highest level of sC/sAC occurrence among the non-conserving amino acids, 40%, compared to less than 25% in the data of the other non-conserving atoms. Moreover, 90% of the observed sC/sAC are anticodons, and not even a single sC/sAC is found in positions 70-72 of tRNA^Thr^.

To further benchmark these statistical results, the occurrence of the triplet CCC was examined. This triplet is expected to be the most prevalent for 14 out of 20 amino acids due to the considerable excess of the nucleotide C in their sequences (Fig. S1). Pre-3’end strings from the tRNAs of the conserving amino acids display CCC considerably less than inferred from their nucleotide distribution. Surprisingly, the strings of Ala, His and Asp, which contain over 44% of nucleotide C in their pre-3’end sequences, hardly show any CCC triplets. Conversely, amino acids whose tRNAs display sC/sACs significantly less than statistically expected (Gln, Tyr, Lys), exhibit the most significant access of CCC.

### A subgroup of six robustly conserving amino acids

The sC/sACs found in the pre-3’end string of the conserving amino acids may appear as a single site which is fully conserved, although the rest of the sequence varies (Pro, Table 1, Fig. 1) or as an accumulation of occurrences from as many as seven different sites (Leu, Val, Fig. 1). The conserved sC/sACs of Ala, Arg, Asp, Gly, His, Pro and Ser are concentrated in one or two sites, each holding mainly one or two coding triplets. In contrast, the overall high occurrence of the conserved sC/sACs of Leu and Val is the sum of small occurrences scattered throughout their pre-3’end strings.

**Figure 1.**
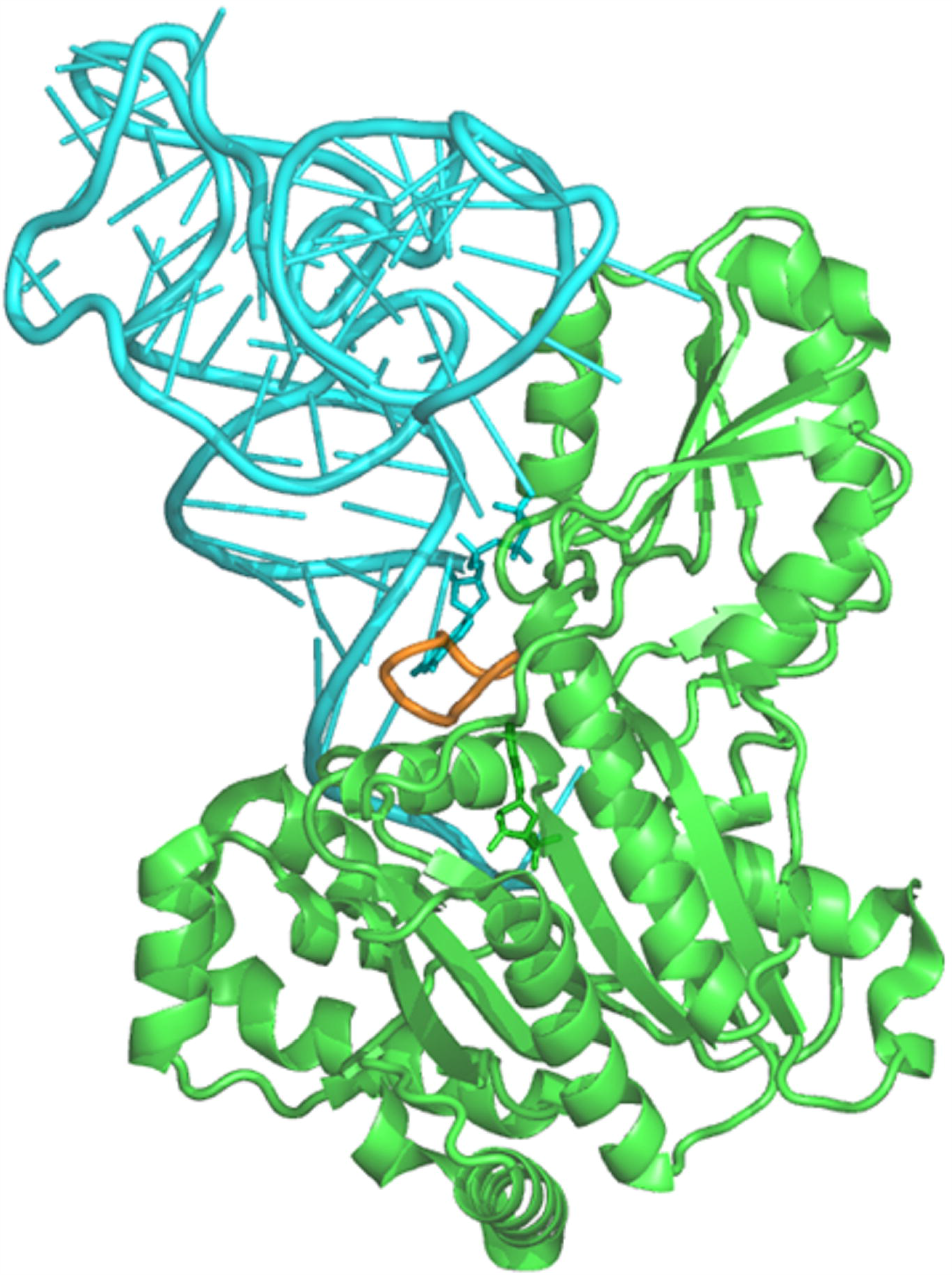
Occurrence of the conserved stem codon and/or anticodon triplets in the pre-3’end string of the conserving amino acids. The column of each “sC/sAC conservation site” is marked by the identity of its conserved triplets and by the percentages of sequences carrying these triplets in the site. * The overall occurrence of the conserved sC/sACs of Ser in its two partially overlapping conservation sites (positions 68-70 and 70-72).

Components of a conserved set are considered as “robustly conserved sC/sACs” when they appear, alone or with one additional coding triplet, at a specific location in more than 60% of the pre-3’end strings belonging to a certain amino acid. This location is termed a “sC/sAC conservation site”. Overall, the occurrence histogram (Fig. 1) contains nine such sites: one for Pro (positions 70-72, 98% occurrence of the codon CCG); Gly (positions 68-70, 94% occurrence of the anticodons CCC or UCC); Asp (positions 64-66, 91% occurrence of the anticodon GUC); Ala (positions 68-70, 64% occurrence of the codon GCU. In the remaining sequences only C69 and U70 are conserved); Arg (positions 66-68, 60% occurrence of the codons CGG or AGG); two sites for Ser (in two partially overlapping positions; 68-70, 63% occurrence of the codons UCU or UCA and positions 70-72, 72% occurrence of UCC, which combine to 94% occurrence of a conserved codon in positions 68-72) and two for His (positions 63-65, 60% occurrence of the codon CAU. In the remaining sequences, only C63 and U65 are conserved. Positions 70-72, 69% occurrence of the codon CAC. In the remaining sequences, only C70 and C72 are conserved, mostly due to the exchange of A71 into G71 in the sequences from *Bacillus*). Interestingly, in contrast to the accumulation of coding triplets in positions 68-70 and 70-72 (Fig. 1), there are hardly any conserved triplets in positions 69-71. The corresponding amino acids and coding sets (Table 2) are termed “robustly conserving amino acids” and “robustly conserved sC/sAC sets” (materials and methods).

In a second benchmark test the occurrences of individual coding triplets per each three-mer location in the pre-3’end strings of every amino acid were counted, without relating to the nucleotide distribution. Cognate sC/sACs were found at a specific location in more than half of the strings belonging to each of the six robustly conserving amino acids, but not in the data of the other 14 amino acids.

Five out of the six robustly conserving amino acids, i.e., Ala, Gly, His, Pro and Ser, have synthetases that belong to class IIa. The statistical correlation between the list of amino acids that robustly conserve the sC/sACs, i.e., Ala, Asp, Gly, His, Pro and Ser, and those whose synthetases belong to class IIa, i.e., Ala, Gly, His, Pro, Ser and Thr [21], was computed. The probability P that such a correlation or higher, could have been obtained randomly, is given by (Materials and Methods):

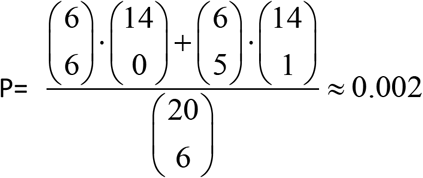

indicating that the null hypothesis of no correlation between robust conservation and belonging to class IIa is wrong.

### Archaea and eukarya

The genomic tRNA sequences of archaea and eukarya were not subjected to the aforementioned analysis due to the difficulty to distinguish pseudo-tRNA gene-sequences, which are prevalent in the database (Sprinzl M., private communication), from the authentic ones. The 3’ end tail, NCCA, used for screening the bacterial data, which excluded 40% of the bacterial sequences, appears only in few archaeal tRNA-genes and is absent in eukaryotes. The more reliable elongator-tRNA sequences are too scarce to ensure any statistical validity. Part of the 63 unique archaeal and 117 unique eukaryotic sequences, belonging to the elongator-tRNAs of the 20 amino acids, do display access of cognate sC/sACs in their pre-3’end strings (Files S2, S3), mostly for the same amino acids classified as conserving in bacteria. However, the position and identity of these cognate coding triplets differ from bacteria, leaving open the question of whether the cognate sC/sACs are conserved in the other realms of life besides bacteria.

## Discussion

### Coding triplets in the acceptor-TΨC stem

In total, over 90% of the bacterial pre-3’end sequences from the nine conserving amino acids contain a conserved codon and/or anticodon triplet (Table 2), although the data probably includes unidentified pseudo-tRNA genes randomly carrying the consensus discriminator base, and possibly sequences from additional species besides the *Mollicutes*, whose tRNAs do not comply with the bacterial pattern of sC/sACs conservation. Coding triplets, which are complementary to the conserved sC/sACs presented here, were previously identified in positions 3-5 of prokaryotic tRNAs of four of the conserving amino acids, i.e., Ala, Asp, Gly and Val [22]. Collectively, these observations point to a highly non-random occurrence of conserved sC/sACs, distributed along the pre-3’end strings of the conserving amino acids.

Nucleotide C is the most abundant (47%) of the nucleotides constituting the 12 robustly conserved sC/sAC triplets (Table 2). The conservation is therefore facilitated by the elevated occurrence of nucleotide C (46%) in the bacterial pre-3’end sequences. Yet, the sC/sACs occurrence in the pre-3’end strings of the conserving amino acids is of consequence with regard to both the expectation values derived from the randomized oligonucleotides constructed with the C-rich “sample” distribution (Table 2) and with the individual nucleotide distributions (Table S1a), validating that the conservation of the cognate stem codons and/or anticodons in the pre-3’end string is a genuine phenomenon. Correspondingly, the triplet CCC, which is expected to be highly frequent due to the significant access of nucleotide C in the pre-3’end of 14 amino acids, is nearly absent in the sequences from the tRNAs of the robustly conserving amino acids, likely due to a negative correlation with the marked occurrence of the conserved sC/sACs (Fig. S1).

Our statistical results are in line with the ensemble attributes of the “operational RNA codes”, determined from a bacterial classification and regression tree, which was inferred from acceptor stem sequences of a large variety of bacterial organisms [23]. The tree exhibited low degree of degeneracy for the same five amino acids that display robustly conserved sC/sACs in their acceptor stem, i.e., Ala, Gly, His, Pro and Ser. In contrast, it presented the highest degree of degeneracy for Arg, Leu and Val, indicating the weakening of the selective constraints on their tRNA acceptor stem identity elements. The clear accumulation of most species corresponding to each of the five robustly conserving amino acids, into a single leaf of the tree, corroborates the conservation of coding information in a specific location of their acceptor stems. Likewise, the high degeneracy obtained for Arg, Leu and Val, agrees with the dispersion of the cognate coding triplets throughout their acceptor-TΨC stem (Fig. 1), found in the present study.

### Contemporary significance of the conserved coding triplet

The unexpected retention of cognate coding triplets at specific locations of the acceptor-TΨC stems of Ala, Asp, Gly, His, Pro and Ser poses the question of the biological significance behind this extreme conservation (Table 2, Table S2).

Five of the robustly conserving amino acids, i.e., Ala, Gly, His, Pro and Ser, exhibit conserved sC/sACs in the nucleotide range 68-72 of the acceptor stem (Fig. 1), which forms a contact interface with synthetase [9]. The tRNAs of all these amino acids are charged by class IIa synthetases, when partition into subclasses is made according to the relevant mode of their binding to the tRNA acceptor stem [21]. The probability that this correlation is random is negligible, 0.2%, suggesting that the high conservation of coding triplets in positions 68-72 is connected with their involvement in the aminoacylation mode applied by their cognate class IIa synthetases. This assertion is further supported by the significant decrease in aminoacylation rate of the full-length tRNA^Pro^ on mutation of **C70** and **G72** [24], from the robustly conserved Pro stem-codon **C**C**G** located in positions 70-72 (Fig. 1).

A plausible aminoacylation mechanism that uses the robustly conserved sC/sACs to discriminate between tRNAs or between their derived minimal substrates, requires specific interaction between the catalytic core of the corresponding synthetases and the conserved sC/sACs. The flexible motif 2 loop from the catalytic core of class II synthetases was found to interact, in a base-specific fashion, with nucleotides from the tRNA acceptor stem. In the tRNA^Ser^:SerRS complex, it protrudes into the major groove down to the fifth base pair, interacting with the nucleotide bases [25], that is, with the same nucleotides that are robustly conserved and that constitute the identity determinants for tRNA^Ser^ recognition [15]. In the tRNA^His^:HisRS complex (Fig. 2) residues R115 and Q117 from motif 2 loop approach the bases of A71 and C72 to 3.7Å and 3.9Å, respectively, where A71 and C72 make part of the His robustly conserved codon CAC, located in positions 70-72 of tRNA^His^ (Fig. 1). This mode of interaction provides a feasible mechanism for the specific recognition of these tRNAs, either through the involvement of the conserved coding triplets in stabilizing a specific conformation identified by the synthetase, or via direct contacts of motif 2 loop with the bases of the conserved coding triplets.

**Figure 2.**
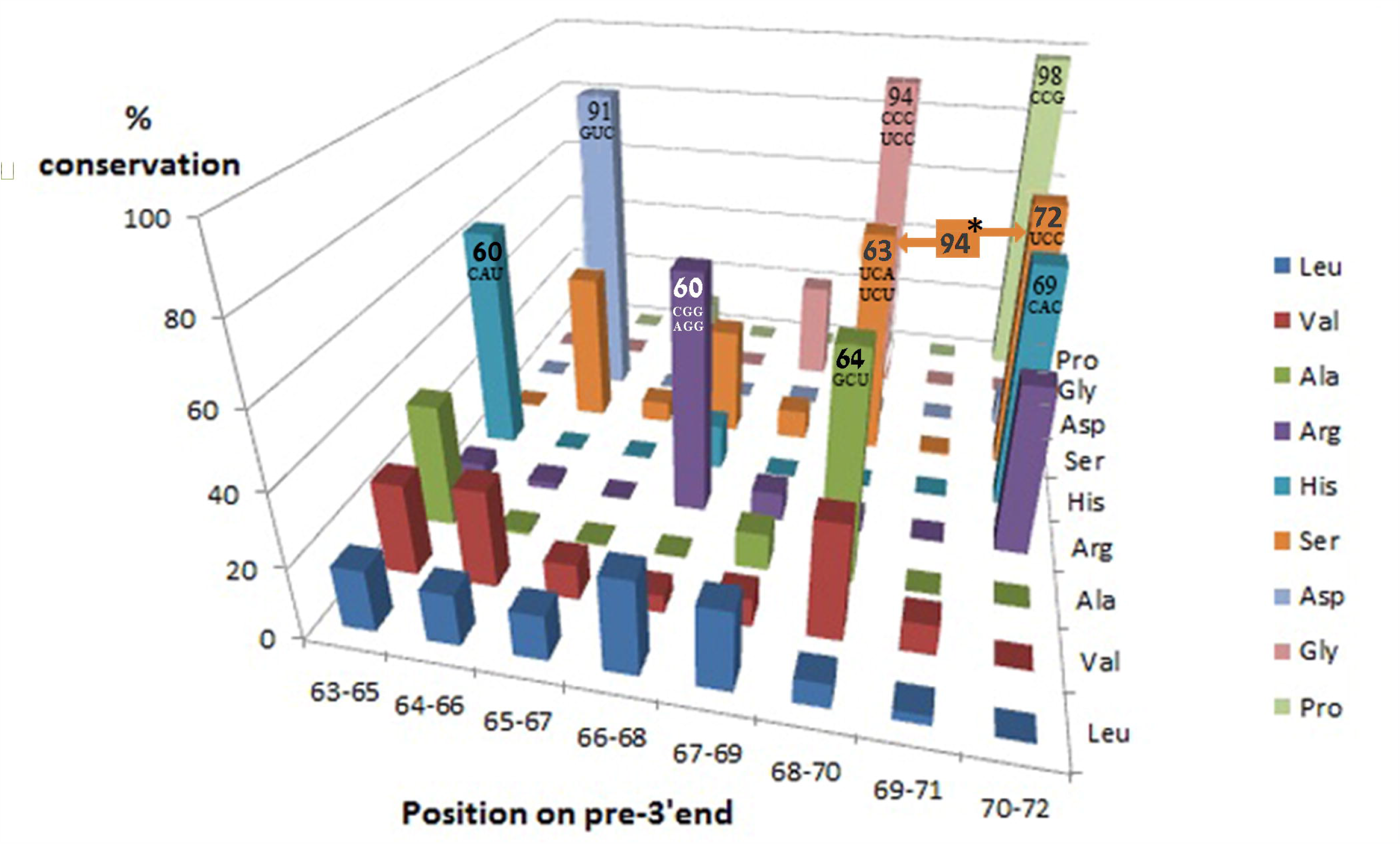
Synthetase interaction with the acceptor stem. tRNA^His^:HisRS complex (pdb 4rdx). Motif 2 loop (orange) of HisRS (green) bulges from the catalytic core into the major groove of tRNA^His^ (cyan) acceptor stem.

Linking the sC/sACs conservation with their being recognized by motif 2 loop, is consistent with the lack of sC/sACs conservation in the pre-3’end strings of the sixth amino acid charged by a class IIa synthetase, Thr (Table S1b). In the structure of the bacterial tRNA^Thr^:ThrRS complex, the identity determinants held in the stem are recognized by the N terminal domain of the enzyme [26] and not by motif 2 loop, a functional exception that can account for the lack of sC/sAC conservation in the stem. Likewise, the partial loss of sC/sAC occurrence in tRNA^Ala^ acceptor stem (Table 2, Fig. 1) can be linked with the reorientation of tRNA^Ala^ relative to the core of its cognate synthetase, as seen in the tRNA^Ala^:AlaRS complex from the archaeon *Archaeoglobus fulgidus* (pdb code 3wqy), which is the only currently available X-ray structure of the complex. Although class II consensus catalytic core exists within this AlaRS structure, motif 2 loop does not interact with the acceptor stem. Recognition of the acceptor stem identity determinants is carried out by the helical loops of α11 and α14, from out of the catalytic core [27], a functional exception that is likely to account for the diminishing sC/sAC conservation in the modern tRNA^Ala^ acceptor stems. The partial conservation of the CAC codon in positions 70-72 of tRNA^His^, which is manifested mainly in the exchange of A71 into G71 in the sequences from the *Bacilli* class, can be possibly linked with a functional variation as well. A second type of synthetase, Histidine tRNA ligase 2, found solely in *Bacillus* [28], may render the sC/sAC conservation in this phylogenetic class surplus.

Involvement of the robustly conserved sC/sACs in the aminoacylation by five class IIa synthetases is further corroborated by the properties of the aminoacylation of minimal RNA substrates. In this process, information contained in the acceptor stem was demonstrated to suffice for charging minihelices derived from the acceptor-TΨC stem, with their cognate amino acids [9,11-16]. This alternative information source, termed the “operational code” [12], depends on identity determinants held in the first five base pairs of the acceptor stem analogs. Aminoacylation of such minihelices was found to be carried out most efficiently for four out of five robustly conserving amino acids i.e., for Ala, Gly, His and Ser [29], while no data is currently available for fifth amino acid, Pro. All but one of the identity determinants recognized by the corresponding synthetases, in the 3’ side of the cognate minihelices, are contained in the robustly conserved coding triplets of these amino acids (Table 3). The coding triplets present in these minihelices can therefore be specifically recognized by motif 2 loop, via the same mechanism suggested for the interaction of the loop with the identity determinants located in acceptor stems of full-length tRNAs. This interaction is likely to facilitates the identification of these minihelices and thus grant them with enhanced efficiency in aminoacylation.

**Table 3.** Identity determinants vs. robustly conserved sC/sACs in the tRNA acceptor stem

| Amino acid |  | **Ala** | **Gly** | **Ser** |  | **His** | **Pro** |
| --- | --- | --- | --- | --- | --- | --- | --- |
| sC/sAC conservation sites in the 68-72 range | Positions | 68-70 | 68-70 | 68-70 | 70-72 | 70-72 | 70-72 |
|  | Conserved sC/sACs* | **GCU** | **CCC/UCC** | **UCU/UCA** | **UCC** | **CAC** | **CCG** |
| Identity determinants for minimal substrate aminoacylation* |  | G3•**U70**<br>[11] | G3:**C70**, C2:G71<sup>^</sup><br>[30] | (C/U)**68**,<br>(C/U)**69**,<br>(A/U)**70**<br>[15] | (A/U)**70**<br>**C71**,<br>**C72**<br>[15] | **C70**,<br>**A71**<br>[30] | **C70\*\***,<br>**G72\*\***<br>[24] |

\* In bold - nucleotides from the robustly conserved coding-triplets serving as identity determinants.
\*\* These identity determinants refer to the recognition of the full-length tRNA.
<sup>^</sup> The identity determinant for aminoacylating minihelices of Gly, C2:G71, is not part of the conserved coding triplets.

The linkage made here between sC/sAC conservation and the contemporary non-AC aminoacylation, can be validated or refuted experimentally. Due to the high occurrence of the conserved sC/sACs in the acceptor stems of the robustly conserving amino acids, the more efficient aminoacylation rates reported in the literature correspond to minihelices carrying a conserved coding triplet. Decrease in aminoacylation rate following the mutation of a non-determinant nucleotide in a robustly conserved coding triplet, can corroborate this linkage.

At the other half of the pre-3’end string, nucleotides 63–67 from the acceptor-TΨC stem establish the contact interface with EF-Tu [3]. The sC/sAC conservation sites in that range, that is, the Asp codon GUC in positions 64-66, which is conserved in 91% of the sequences, and the His codon CAU in positions 63-65, which is conserved in 60% of the sequences (Fig. 1), are suggested to contribute to the binding affinity for EF-Tu. In particular, EF-Tu was shown to use the tRNA^Asp^ base pairs G51:C63, C50:**G64** and G49:**U65** for tightly binding it [31], where **G64** and **U65** are included in the conserved Asp codon GUC. For His, only C63 and U65 are completely conserved, suggesting that the present involvement of specific nucleotides in the recognition of tRNA by EF-Tu is likely to determine their degree of conservation.

The sC/sACs conserved in pre-3’end strings of the non-robustly conserving amino acids Arg, Leu and Val, which are presently charged by class I synthetases, seem to lack a contemporary role. The conserved sC/sACs of Leu and Val are dispersed throughout the pre-3’end string (Fig. 1) rather than holding specific sites, as would be expected from functional sC/sACs. Consistently, the identity determinants for aminoacylating minimal analogs of tRNA^Leu^ are found in the D-arm and in A73, but not in the pre-3’end [32]. 60% of the pre-3’end sequences from tRNA^Arg^ actually contain a robustly conserved codon in positions 66-68 (Fig. 1). However, this site is unlikely to serve as an identity determinant, because its location partially overlaps the ranges of tRNA interaction with EF-Tu and with the synthetase. In accord, the base of A20 in the D-loop is the dominant identity element for non-AC aminoacylation [33].

As expected from the assumed lack of contemporary role, the level of sC/sACs conservation in the pre-3’end strings of these three amino acids is lower compared to the six robustly-conserving amino acids (Table 2). Also, the number of conserved sC/sACs per amino acid is larger; five, six and eight for Leu, Arg and Val, compared with one to three for the robustly conserving amino acids. Moreover, the robustly conserved sC/sAC sets, which comprise just codon or just anticodons, are evolutionary more plausible, because they can be the descendants of a single cognate coding triplet that existed in the acceptor-TΨC stem of the ancestral tRNA, while those containing both codons and anticodons, as is the case with Leu, Arg and Val, cannot. In accord, the “operational RNA codes” inferred from the bacterial classification and regression tree, demonstrated the highest degree of degeneracy for Arg, Leu and Val, indicating the reduced conservation of their acceptor stem identity elements [23]. These statistical observations seem to point to a gradual process by which, in the absence of a contemporary role, the conserved sC/sACs of Leu, Arg and Val, were shifted throughout the acceptor-TΨC stem, while random mutations supplemented the list of conserved sC/sACs and lowered the overall level of conservation.

Taken together, these five amino acids, i.e. Ala, Gly, His, Pro and Ser, share three different acceptor-stem related features: robust conservation of the sC/sAC, enhanced efficiency of minimal-substrates aminoacylation, and being charged by class IIa synthetases. This correlation suggests the existence of a variant of the aminoacylation mode of action applied by the corresponding class IIa synthetases, which takes advantage of the information embedded in the robustly conserved sC/sACs to discriminate between tRNAs. Such a mechanism can provide reasoning for the enhanced efficiency of charging these minihelices and more importantly - for the extreme conservation (Table 2, Table S2) of the coding triplets in the modern acceptor stems of these amino acids.

### Possible evolutionary implication of the conserved coding triplet

The level of sC/sAC occurrence in the acceptor-TΨC arm is highly correlated with the antiquity of the corresponding amino acids. Eight out of the nine conserving amino acids (Table 2), i.e., all except His, are listed among the 10 earliest appearing amino acids, according to a consensus chronology built on 60 criteria [34], which is compatible with the results of the Miller-Urey experiment [35]. In particular, the robustly conserving amino acids Ala, Asp, Gly, Pro and Ser are among the six most ancient amino acids. The probability of such a correlation being obtained at random is 0.002, suggesting that the origin of the coding triplets, conserved in the acceptor-TΨC stem, is prebiotic [22].

The evolutionary history of tRNA recognition has gained considerable interest over the years, due to its linkage with the origin of the genetic code, eliciting a plethora of hypotheses. The capability of modern synthetases to utilize information held in the first base pairs of the acceptor stem for specific aminoacylation, gave rise to the conjecture that this mode of operation is ancestral. The limited size of the primordial synthetase permitted it to reach only recognition elements residing in the acceptor stem, in the vicinity of the amino acid attachment site [36,37]. This simple proto-synthetase, which is believed to correspond to the modern catalytic domain [12,14], was suggested to have activated amino acids and to specifically recognize the tRNA via motif 2 loop (Fig. 2), charging in this manner early tRNAs on the basis of their acceptor stem identity determinants [38]. Rodin et al. [39] further proposed that the operational RNA code as we see it today, actually developed from a few ancient anticodon-codon-like pairs located in the first positions of the acceptor stem, and that unrecognized relationship between acceptor and anticodon points to the historically common root of the two codes.

In line with these hypotheses, we suggest that the primordial aminoacylation of proto-tRNAs by their cognate class IIa synthetases was controlled by the full coding triplets, which are still robustly conserved in the acceptor stem, rather than merely by the discrete identity determinants. The existence of a feasible non-AC aminoacylation mechanism that relies on the sC/sACs, which are conserved only in the acceptor stems of Ala, Gly, His, Pro and Ser, points to ancestry of class IIa synthetases, in accord with previous proposals [40,41]. The robustly conserved coding triplets, suggested here to control the prebiotic aminoacylation, form a subset of the standard genetic code. They incorporate all but one of the cognate identity determinants that constitute the “second genetic code” (Table 3), suggesting that the “second code” [12,17], which was proposed to be distinct from the standard genetic code [16], actually makes part of it. In accord with the conjecture concerned with the common root of the acceptor and anticodon codes [39], the codons and anticodons conserved in the acceptor-TΨC stem are suggested here to have had a primal role in establishing the extant genetic code.

The three key components of the protein-synthesis machinery, i.e., the tRNA, synthetase and EF-Tu, along with the ribosome, have coevolved into the intricate contemporary translation system, still keeping traces of their prebiotic roots in the form of the conserved coding-triplets. The evolutionary path proposed by the coding triplets conserved in the tRNA acceptor stem, commencing from specificity control in a primordial aminoacylation, to the involvement in the specificity recognition by the contemporary synthetase, outlines a continuous route from the prebiotic era to modern biology.

## Conclusions

The extreme conservation of cognate coding triplets in the acceptor stem of five amino acids having tRNAs charged by class IIa synthetases, points to their involvement in an extant function. A contemporary aminoacylation mode, whereby the coding triplets conserved in these tRNA acceptor stems, or in minihelices derived from them, assist the recognition by their cognate synthetases, is likely to account for it. The antiquity of the corresponding amino acids suggests that this mode of action is a vestige of a simple ancestral process of aminoacylating proto-tRNAs via the acceptor stem coding triplets. This, in turn, suggests the ancestry of class IIa synthetases, the incorporation of the “second genetic code” within the standard code and the involvement of the stem coding triplets in the establishment of the standard genetic code.

## Materials and Methods

### Dataset construction

Sequences of the 3’ side of the acceptor-TΨC stem from the bacterial tRNAs of each of the 20 canonical amino acids, were obtained from the tRNAdb database [19,20], http://trna.bioinf.uni-leipzig.de, 2018. The tRNAdb database holds over 6200 bacterial tRNA genes, belonging to the 20 canonical amino acids. Only tRNA sequences having a full CCA 3’ end and a consensus discriminator base (N73) [10] were incorporated into the statistics, with one exception; the data for Glu contained similar amount of sequences with the canonical G73 and with the non-canonical A73, both groups displaying the same pattern of sC/sAC occurrence. To obtain as much data as possible, tRNAs with non-canonical A73 were incorporated as well. After the exclusion of the partial sequences and of the rest of the non-canonical data, the initial data set used in the current statistical analysis contained 60% of the bacterial tRNA genes from the database.

The observed occurrence of the cognate codon and anticodon triplets was determined in 10-mer strings consisted of nucleotides 63-72 (termed pre-3’end), by scanning the sequences. Nucleotides 61 and 62 were not included in the study, being conserved and semi-conserved, respectively. The scanning indicated that one class of bacteria, i.e., species from six genera of the *Mollicutes* class, which constituted about 9% of the initial data set (350 gene-sequences), does not share the statistical behavior of the other bacteria (Table 1, File S1, Table S2) and it was excluded. No other species in the data contained enough entries to be reliably recognized as diverging from the conservation pattern. Excluding the *Mollicutes* resulted in a final data set consisting of 3324 tRNA gene-sequences from over 100 bacterial species.

### Definitions

#### Conserving amino acid

Amino acids for which more than 85% of their pre-3’end strings contain cognate sC/sACs. Amino acids for which less than 40% of their pre-3’end strings contain cognate sC/sACs are denoted “non-conserving amino acids”.

#### Conserved sC/sAC set

For the conserving amino acids, is the minimal set of cognate triplets accounting for 98% of the sequences carrying a cognate coding triplet.

#### Robustly conserved sC/sACs

are conserved sC/sACs that are located at specific sites in the pre-3’end string, occurring, alone or with a single additional coding triplet (which is also considered as robustly conserved), in more than 60% of the sequences.

#### Robustly conserved sC/sAC set

of a specific amino acid, is the set containing all its robustly conserved sC/sACs, conditional on the requirement that the sC/sAC set is composed of either just codons or just anticodons.

#### Robustly conserving amino acids

Are those having a robustly conserved sC/sAC set.

### Calculation of expected occurrences

The expected occurrence of cognate coding triplets in the pre-3’end string, was determined by analyzing their occurrence in simulated 10-mer randomized oligonucleotides. A pool of 10^5^ randomized strings was generated using MATLAB and Statistics Toolbox (Release 2012b, The MathWorks, Inc., Natick, Massachusetts), by producing a string of 10^6^ nucleotides with a predetermined nucleotide-distribution, randomizing it and cutting it into 10^5^ 10-mer oligonucleotides. For the conserving amino acids, the percentage of the 10-mer randomized strings carrying at least one cognate triplet from the conserved triplet-set is defined as the expected occurrence. For the remaining amino acids, for which a conserved set of sC/sACs does not exist, the percentage of the 10-mer randomized strings carrying at least one cognate codon or anticodon triplet was set to be the expected occurrence. The error value for the expected percentage, ±0.8%, includes a statistical error of 0.6% and a rounding error of 0.5%. The probability of random occurrence, *P*, was calculated for each amino acid, from the results of a χ^2^ test, using a statistical cutoff of *P* ≤ 0.01.

### Benchmark test for robust conservation

The test counts the occurrences of all (coding and non-coding) triplets according to their appearance per 3-mer site within the 10-mer pre-3’end strings. Any cognate coding triplet that appears alone at a specific location, in most of the strings of an amino acid, is assumed to denote a highly conserving amino acid.

### Calculation of correlation

The probability of obtaining exactly m common objects in two groups of n1, n2 objects chosen randomly out of a batch of n objects, is given by:

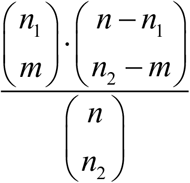

where:

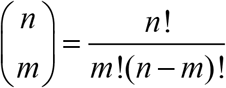

## Acknowledgements

We thank D. Agmon for generating the MATLAB program, the late A. Cohen for the statistical consultancy and S. Agmon for his assistance.

## Abbreviations

AC: anticodon
sC/sAC: stem-codon and/or stem-anticodon
aa: amino acid
aaRS: aminoacyl-tRNA synthetase
pre-3’end: a string of ten nucleotides preceding the tRNA 3’ end

---

**Fig S1:**
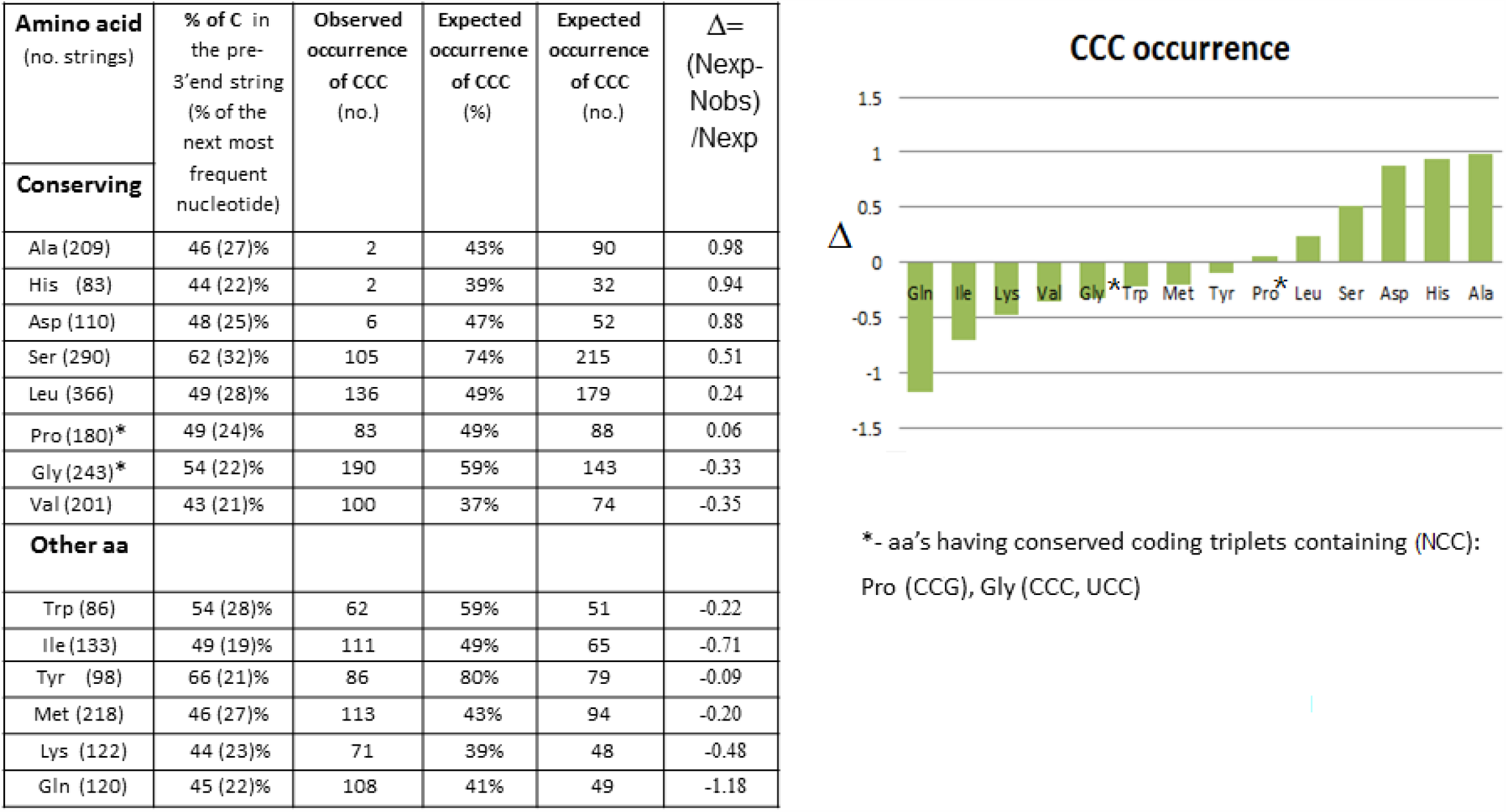
Benchmark-Observed pre3’-end sequences containing CCC in amino acids with large excess on nucleotide C. The occurrence of the triplet CCC is expected to be the most prevalent in the 14 amino acids shown in the figure, due a significant over-representation of C nucleotides in their pre-3’end (access of 19-44% compared to the next most frequent nucleotide). However, strings from the statistically most conserving amino acids, i.e. Ala, His, Asp, Ser, and Leu, display pre-3’end containing CCC considerably less than inferred from their nucleotide distribution and surprisingly, the first three amino acids, i.e. Ala, His and Asp, hardly show any CCC triplets in their strings (2, 2 and 6, respectively). The highly conserving amino acids Pro and Gly, conserve (CCG) and (CCC, UCC), respectively, which is manifested in the expected increase in CCC occurrence in their strings, relative to the other conserving amino acids. At the other end of the scale, the least C/AC-exhibiting amino acid, Gln, displays the most significant access of CCC.

