## Supplementary material for "Coding triplets in the tRNA acceptor-TΨC arm and their role in present and past tRNA recognition": DataSet

**DataSet S1: Bacteria- occurrence of coding triplet in the pre-3'end string.** All the unique pre-3'end strings (nucleotides 63-72) of bacterial elongator-tRNAs (not genes) sequences from the tRNAdb[1,2], http://trna.bioinf.uni-leipzig.de, belonging to the 20 amino acids, clustered according to their conservation type determined from the bacterial tRNA gene-sequences. Conserved codons are framed and conserved anticodons underlined. Mycoplasma (Mollicutes) sequences are grouped at the end for each amino acid.

### Bacteria- Conserving amino acids:

#### Ala

63-72

```
>tdbR00000007|Bacillus_subtilis|1423|Ala|5GC
-GGAGCCUUAGCUCAGCD-GGG--AGAGCGCCUGCUU5GC=CGCAGGAG-------------------7UCAGCGGTPCGAUCCCGCUAGGCUCCACCA
>tdbR00000008|Escherichia_coli|562|Ala|GGC
-GGGGCUANAGCUCAGCD-GGG--AGAGCGCUUGCAUGGCAUGCAAGAG-------------------7UCAGCGGTPCGAUCCCGCUUAGCUCCACCA
>tdbR00000009|Escherichia_coli|562|Ala|VGC
-GGGGGCA4AGCUCAGCD-GGG--AGAGCGCCUGCUUVGCACGCAGGAG-------------------7UCUGCGGTPCGAUCCCGCGCGCUCCCACCA
>tdbR00000010|Escherichia_coli|562|Ala|VGC
-GGGGCUAUAGCUCAGCD-GGG--AGAGCGCCUGCUUVGCACGCAGGAG-------------------7UCUGCGGTPCGAUCCCGCAUAGCUCCACCA

>tdbR00000005|Mycoplasma_capricolum|2095|Ala|UGC
-GGGCCCUNAGCUCAGCD-GGG--AGAGCACCUGCCUUGC=CGCAGGGG-------------------7UCGACGGUPCGAUCCCGUUAGGGUCCACCA
=================================================
```

#### Arg

```
>tdbR00000354|Bacillus_subtilis|1423|Arg|ICG
-GCGCCCGUAGCUCAAU--GGAD-AGAGCGUUUGACUICGKAUCAAAAG-------------------7UUAGGGGTPCGACUCCCCUCGGGCGCGCCA
>tdbR00000359|Escherichia_coli|562|Arg|CCG
-GCGCCCGUAGCUCAGCD-GGAD-AGAGCGCUGCC%UCCGKAGGCAGAG-------------------7UCUCAGGTPCGAAUCCUGUCGGGCGCGCCA
>tdbR00000355|Escherichia_coli|562|Arg|ICG
-GCAUCCG4AGCUCAGCD-GGD--AGAGUACUCGG%UICG/ACCGAGCG-------------------7XCGGAGGTPCGAAUCCUCCCGGAUGCACCA
>tdbR00000357|Escherichia_coli|562|Arg|{CU
-GUCCUCUUAGUUAAAU--GGAD-AUAACGAGCCC%U{CU6AGGGCUAA-------------------U-UGCAGGTPCGAUUCCUGCAGGGGACACCA
>tdbR00000358|Escherichia_coli|562|Arg|{CU
-GCGCCCUUAGCUCAGUU-GGAU-AGAGCAACGAC%U{CU6AGPCGUGG-------------------GCCGCAGGTPCGAAUCCUGCAGGGCGCGCCA

>tdbR00000353|Mycoplasma_capricolum|2095|Arg|!CU
-GCCCAUGUAGCUCAGUA-GGAD-AGAGCACGCGCCU!CU6AGCGUGAG-------------------7UCGGAAGUPCGAGCCUUCUCGUGGGCACCA
>tdbR00000352|Mycoplasma_capricolum|2095|Arg|ICG
-GCGCCCGNAGAUCAAUD-GGAD-AGAUCGCUUGACUICGKAPCAAAAG-------------------7UUGGGGGUPCGAGUCCCUCCGGGCGCACCA
=================================================
```

#### Asp

```
>tdbR00000025|Thermus_thermophilus|274|Asp|GUC
-GGCCCCG4GGUGPAGUU-#GDD-AACACACCCGCCUGUCACGPGGGAG-------------------AUCGCGGGFPCG"GUCCCGUCGGGGCCGCCA

>tdbR00000024|Mycoplasma_capricolum|2095|Asp|GUC
-GGCCCCANAGCGAAGUD-GGDD-AUCGCGCCUCCCUGUCACGGAGGAG-------------------AUCACGGGUPCGAGUCCCGUUGGGGUCGCCA
=================================================
```

#### Gly

```
>tdbR00000116|Bacillus_subtilis|1423|Gly|!CC
-GCGGGUGUAGUUUAGU--GGD--AAAACCUCAGCCU!CCAAGCUGAUG-------------------U-CGUGAGTPCGAUUCUCAUCACCCGCUCCA
>tdbR00000117|Escherichia_coli|562|Gly|CCC
-GCGGGCG4AGUUCAAU--GGD--AGAACGAGAGCUUCCCAAGCUCUAU-------------------A-CGAGGGTPCGAUUCCCUUCGCCCGCUCCA
>tdbR00000118|Escherichia_coli|562|Gly|GCC
-GCGGGAAUAGCUCAGDD-GGD--AGAGCACGACCUUGCCAAGGUCGGG-------------------7UCGCGAGTPCGAGUCUCGUUUCCCGCUCCA
>tdbR00000121|Salmonella_typhimurium|602|Gly|NCC
-GCGGGCAUCGUAUAAU--GGCU-AUUACCUCAGCCUNCCAAGCUGAUG-------------------A-UGCGGGTPCGAUUCCCGCUGCCCGCUCCA
>tdbR00000114|Staphylococcus_epidermidis|1282|Gly|UCC
-GCGGGAG4AUUUCAACU-UUD--AGAAUACGUUCCUUCCCGGAACGAG-------------------A-UAUAGGUGCAAAUCCUAUCUUCCGCUCCA
>tdbR00000113|Streptomyces_coelicolor_A3(2)|100226|Gly|CCC
-GCGGGUGUAGUUCAAU--GGD--AGAACAUCAGCUUCCCAAGCUGAGA-------------------G-CGCGAGTPCGAUUCUCGUCACCCGCUCCA

>tdbR00000111|Mycoplasma_capricolum|2095|Gly|UCC
-GCAGGUGNAGUUUAAU--GGD--AGAACUUCAGCCUUCC=AGCUGAUU-------------------G-UGAGGGUPCGAUUCCCUUCACCUGCUCCA
=================================================
```

#### His

```
>tdbR00000141|Salmonella_typhimurium|602|His|QUG
GGUGGCUA4AGCUCAGDD-GGD--AGAGCCCUGGAUUQUG/PPCCAGUU-------------------7UCGUGGGTPCGAAUCCCAUUAGCCACCCCA

>tdbR00000139|Mycoplasma_capricolum|2095|His|GUK
GGCGUAGGUGGUGAAGU--GGDD-A"CACAUCAGGUUGUKKCPCUGACA-------------------UACGCGGGUPCGAUCCCCGUUCUACGCCCCA
```

63-72

#### Leu

```
>tdbR00000219|Bacillus_subtilis|1423|Leu|CAG
-GCGGGUGUGCGGGAAUD-GGDA-GACCGGCUAGAUUCAGKAPCUAGG-GUCU---UUAU----GGACC-UGAGGGTPCA"GUCCCUUCACCCGCACCA
>tdbR00000595|Escherichia_coli|562|Leu|BAA
-GCCGAAG4GGCGAAADC-#GDA-GACACAGUUGAPUBAA*APCAACC-GUA----GAAA-----UACG-UGCCGGTPCGAGUCCGGCCUUCGGCACCA
>tdbR00000222|Escherichia_coli|562|Leu|GAG
-GCCGAGGUGGUGGAADD-#GDA-GACACGCUACCUUGAG;PGGUAGU-GCCC---AAUA----GGGCU-UACGGGTPCAAGUCCCGUCCUCGGUACCA
>tdbR00000220|Escherichia_coli|562|Leu|HAA
-GCCCGGA4GGUGGAADC-#GDA-GACACAAGGGAPUHAA*APCCCUC-GGCG---UUCG----CGCUG-UGCGGGTPCAAGUCCCGCUCCGGGUACCA
>tdbR00000227|Geobacillus_stearothermophilus|1422|Leu|BAA
-GCCGAUG4GGCGGAAUD-GGCA-G"CGCGCACGACUBAA*APCGUGU-GGGC---UUU-----GCCCG-UGUGGGTPCGACUCCCACCAUCGGCACCA
>tdbR00000224|Rhodospirillum_rubrum|1085|Leu|CAA
-GCCUUUGUAGCGGAAD--GGD--AACGCGGCAGACUCAAHAPCUGCU-UUGG---UAAC----CCAGG-UGGUAGTPCGACUCUCCCCAAAGGCACCA
>tdbR00000223|Salmonella_typhimurium|602|Leu|CAG
-GCGAAGGUGGCGGAADD-#GDA-GACGCGCUAGCUUCAG;PGPUAGU-GUCC---UUAC----GGACG-UGGGGGTPCAAGUCCCCCCCUCGCACCA
>tdbR00000225|Synechococcus_elongatus_PCC_6301|269084|Leu|CAA
-GGGCAAGUGGCGGAAUD-GGDA-GACGCAGCAGACUCAAHAPCUGCC-GCUA---GCGA----UAGUG-UGUGGGTPCGAGUCCCACCUUGCCCACCA
>tdbR00000226|Synechococcus_elongatus_PCC_6301|269084|Leu|CAG
-GCGGAACUGGCGGAAUD-GGDA-GACGCGCUAGAUUCAGKPPCUAGU-GGUU---UCAC----GACUG-UCCGGGTPCAAGUCCCGGGUUCCGCACCA

>tdbR00000217|Mycoplasma_capricolum|2095|Leu|)AA
-CCCCAAGNGGCGGAAUA-GGDA-G"CGCAUUGGACU)AA=APCCAAC-GGGC---UUAAU---AUCCUGUGCCGGUPCAAGUCCGGCCUUGGGGACCA
>tdbR00000216|Mycoplasma_capricolum|2095|Leu|BAA
-GCCUUUUUGGCGGAAUD-GGCA-G"CGCAUUAGACUBAA=APCUAAC-GAA----GAAA-----UUCG-UAUCGGUPCGAAUCCGAUAAAGGGCACCA
>tdbR00000218|Mycoplasma_capricolum|2095|Leu|UAG
-GGGGGAUNGGCGGAAUD-GGCA-G"CGCACUAGACUUAGKAPCUAGC-GUC----UUU------GACG-UAAGGGUPCAAGUCCCUUAUCCCCCACCA
=================================================
```

#### Pro

```
>tdbR00000316|Bacillus_subtilis|1423|Pro|5GG
-CGGGAAGUAGCUCAGCUUGGD--AGAGCACAUGGPU5GGKACCAUGGG-------------------7UCGCAGGTPCGAAUCCUGUCUUCCCGACCA
>tdbR00000623|Escherichia_coli|562|Pro|CGG
-CGGUGAU4GGCGCAGCCUGGD--AGCGCACUUCGJUCGGKACGAAGGG-------------------7UCGGAGGTPCGAAUCCUCUAUCACCGACCA
>tdbR00000318|Salmonella_typhimurium|602|Pro|GGG
-CGGCACG4AGCGCAGCCUGGD--AGCGCACCGUCBUGGGKUPGCGGGG-------------------7UCGGAGGTPCAAAUCCUCUCGUGCCGACCA

>tdbR00000314|Mycoplasma_capricolum|2095|Pro|UGG
-CGGGAAGUGGCUCAGUUUGGD--AGAGCAUUCGGUUUGGKACCGAAGG-------------------7NCGCAGGUPCAAAUCCUGUCUUCCCGACCA
=================================================
```

#### Ser

```
>tdbR00000387|Bacillus_subtilis|1423|Ser|5GA
-GGAGGAA4ACCCAAGUCUGGCDGA"GGGAUCGGUCU5GA+AACCGACAGGGUG--UCAA---AGCCCG-CGGGGGTPCGAAUCCCUCUUCCUCCGCCA
>tdbR00000388|Bacillus_subtilis|1423|Ser|GCU
-GGAGAAGUACUCAAGU--GGCDGA"GAGGCGCCCCUGCU6AGGGUGU_GUCGC--GUAA---GCGGCG-CGAGGGTPCAAAUCCCUCCUUCUCCGCCA
>tdbR00000389|Bacillus_subtilis|1423|Ser|GGA
-GGAGAGC4GUCCGAGU--GGDCGA"GGAGCACGAUUGGAAAPCGUGUAGGCGGU-CAAC--UCCGUCU-CAAGGGTPCGAAUCCCUUGCUCUCCGCCA
>tdbR00000390|Escherichia_coli|562|Ser|CGA
-GGAGAGAUGCCGGAGC--#GCDGAACGGACCGGUCUCGA*AACCGGA-GUAGGG-GCAA--CUCUAC--CGGGGGTPCAAAUCCCCCUCUCUCCGCCA
>tdbR00000391|Escherichia_coli|562|Ser|GCU
-GGUGAGG4GGCCGAGA--GGCDGAAGGCGCUCCC%UGCU6AGGGAGU-AUGCGGUCAAAAGCUGCAU--CCGGGGTPCGAAUCCCCGCCUCACCGCCA
>tdbR00000393|Escherichia_coli|562|Ser|GGA
-GGUGAGG4GUCCGAGU--#GDDGAAGGAGCACGCCUGGAAAGPGUGU-AUACG--GCAA---CGUAU--CGGGGGTPCGAAUCCCCCCCUCACCGCCA
>tdbR00000394|Escherichia_coli|562|Ser|VGA
-GGAAGUG4GGCCGAGC--#GDDGAAGGCACCGGUBUVGA*AACCGGC-GACCC--GAAA---GGGUU--CCAGAGTPCGAAUCUCUGCGCUUCCGCCA

>tdbR00000384|Mycoplasma_capricolum|2095|Ser|GCU
-GGGUUAANACUCAAGUD-GGDG-A"GAGGACACCCUGCU6AGGUGUUAGGUCGG-UCU---CCGGCG--CGAGGGUPCGAGUCCCUCUUAACCCGCCA
>tdbR00000385|Mycoplasma_capricolum|2095|Ser|UGA
-GGAAGAUNACCCAAGUCCGGCDGA"GGGAUCGGUCUUGA=AACCGAGAGUCGG--GGAAA--CCGAG--CGGGGGUPCGAAUCCCUCAUCUUCCGCCA
```

#### Val

```
>tdbR00000453|Bacillus_subtilis|1423|Val|5AC
-GGAGGAUUAGCUCAGCD-GGG--AGAGCAUCUGCCU5AC=AGCAGAGG-------------------7UCGGCGGTPCGAGCCCGUCAUCCUCCACCA
>tdbR00000454|Escherichia_coli|562|Val|GAC
-GCGUCCG4AGCUCAGDD-GGDD-AGAGCACCACCUUGACAUGGUGGGG-------------------7XCGGUGGTPCGAGUCCACUCGGACGCACCA
>tdbR00000455|Escherichia_coli|562|Val|GAC
-GCGUUCA4AGCUCAGDD-GGDD-AGAGCACCACCUUGACAUGGUGGGG-------------------7XCGUUGGTPCGAGUCCAAUUGAACGCACCA
>tdbR00000456|Escherichia_coli|562|Val|VAC
-GGGUGAU4AGCUCAGCD-GGG--AGAGCACCUCCCUVAC=AGGAGGGG-------------------7UCGGCGGTPCGAUCCCGUCAUCACCCACCA
>tdbR00000457|Geobacillus_stearothermophilus|1422|Val|GAC
-GAUUCCGUAGCUCAGCD-GGG--AGAGCGCCACCUUGAC=GGGUGGAG-------------------7UCGCUGGTPCGAGCCCAGUCGGAAUCACCA

>tdbR00000451|Mycoplasma_capricolum|2095|Val|UAC
-GGAGUGUUAGCUCAGCD-GGG--AGAGCUCCUGCCUUAC=AGCAGGCG-------------------7UCAUAGGUPCAAGUCCUAUACACUCCACCA
=================================================
```

### Bacteria- non conserving amino acids

#### Gln

63-72

```
>tdbR00000332|Escherichia_coli|562|Gln|CUG
-UGGGGUA4CGCCAAGC--#GD--AAGGCACCGGAJUCUG/PPCCGGCA-------------------UUCCGAGGTPCGAAUCCUCGUACCCCAGCCA
>tdbR00000333|Escherichia_coli|562|Gln|NUG
-UGGGGUA4CGCCAAGC--#GD--AAGGCACCGGUJUNUG/PACCGGCA-------------------UUCCCUGGTPCGAAUCCAGGUACCCCAGCCA
=================================================
```

#### Glu

```
>tdbR00000045|Escherichia_coli|562|Glu|SUC
-GUCCCCUUCGUCPAGA--GGCCCAGGACACCGCCCUSUC/CGGCGGUA-------------------A-CAGGGGTPCGAAUCCCCUGGGGGACGCCA
>tdbR00000046|Escherichia_coli|562|Glu|SUC
-GUCCCCUUCGUCPAGA--GGCCCAGGACACCGCCCUSUC/CGGCGGUA-------------------A-CAGGGGTPCGAAUCCCCUAGGGGACGCCA
>tdbR00000048|Synechocystis_sp.|1143|Glu|NUC
-GCCCCCAUCGUCUAGA--GGCCDAGGACACCUCCCUNUCACGGAGGCG-------------------A-CAGGGATPCGAAUUCCCUUGGGGGUACCA

>tdbR00000044|Mycoplasma_capricolum|2095|Glu|!UC
-GGCCUGUUGGUGAAGC--GGDD-A"CACACACGGUU!UCAUCCGUGGA-------------------CACACGGGUPCGAACCCCGUACAGGCUACCA
=================================================
```

#### Lys

```
>tdbR00000179|Bacillus_subtilis|1423|Lys|$UU
-GAGCCAUUAGCUCAGUD-GGD--AGAGCAUCUGACU$UUHAPCAGAGG-------------------7UCGAAGGTPCGAGUCCUUCAUGGCUCACCA
>tdbR00000181|Escherichia_coli|562|Lys|SUU
-GGGUCGUUAGCUCAGDD-GGD--AGAGCAGUUGACUSUU6APCAAUUG-------------------7XCGCAGGTPCGAAUCCUGCACGACCCACCA

>tdbR00000178|Mycoplasma_capricolum|2095|Lys|!UU
-GACUCGUUAGCUCAGCC-GGD--AGAGCAACUGGCU!UU6ACCAGUGG-------------------7UCCGGGGUPCGAAUCCCCGACGAGUCACCA
>tdbR00000177|Mycoplasma_capricolum|2095|Lys|CUU
-GUCUGAUUAGCGCAACD-GGC--AGAGCAACUGACUCUU6APCAGUGG-------------------7UUGUGGGUPCGAUUCCCACAUCAGGCACCA
=================================================
```

#### Met

```
>tdbR00000274|Bacillus_subtilis|1423|Met|CAU
-GGCGGUGUAGCUCAGC--GGCD-AGAGCGUACGGUUCAU=CCCGUGAG-------------------7DCGGGGGTPCGAUCCCCUCCGCCGCUACCA
>tdbR00000276|Escherichia_coli|562|Met|MAU
-GGCUACG4AGCUCAGDD-#GDD-AGAGCACAUCACUMAU6APGAUGGG-------------------7XCACAGGTPCGAAUCCCGUCGUAGCCACCA
>tdbR00000275|Thermus_thermophilus|274|Met|CAU
-CGCGGGG4GGAGCAGCCU#GD--AGCUCGUCGGGBUCAUAACCCGAAG-------------------7UCGCGGGFPCA"AUCCCGCCCCCGCAACCA

>tdbR00000273|Mycoplasma_capricolum|2095|Met|CAU
-GGCGGGGNAGCUCAGUD-GGDD-AGAGCGUUCGGUUCAUACCCGAAAG-------------------7UCGAGAGUPCAAAUCUCUCCCCCGCUACCA
=================================================
```

#### Tyr

```
>tdbR00000543|Bacillus_subtilis|1423|Tyr|QUA
-GGAGGGG4AGCGAAGU--GGCUAA"CGCGGCGGACUQUA+APCCGCU-CCC----UCA------GGGUUCGGCAGTPCGAAUCUGCCCCCCUCCACCA
>tdbR00000545|Escherichia_coli|562|Tyr|QUA
-GGUGGGG4UCCCGAGC--#GCCAAAGGGAGCAGACUQUA*APCUGCC-GUC----AUC------GACUUCGAAGGTPCGAAUCCUUCCCCCACCACCA
>tdbR00000547|Geobacillus_stearothermophilus|1422|Tyr|QUA
-GGAGGGG4AGCGAAGU--#GCUAA"CGCGGCGGACUQUA*APCCGCU-CCC----UUU------GGGUUCGGCGGTPCGAAUCCGUCCCCCUCCACCA

>tdbR00000542|Mycoplasma_capricolum|2095|Tyr|GUA
-GGAGGGGUAGCGAAGU--GGCDAA"CGCGGUGGCUGUA=CCCACUU-CC-----UUAC------GGUUCGGGGGUPCGAAUCCCUCCCCCUCCACCA
==================================================
```

#### Asn

```
>tdbR00000295|Escherichia_coli|562|Asn|QUU
-UCCUCUG4AGUUCAGDC-GGD--AGAACGGCGGACUQUU6APCCGUAU-------------------7UCACUGGTPCGAGUCCAGUCAGAGGAGCCA

>tdbR00000294|Mycoplasma_capricolum|2095|Asn|GUU
-GGCUUUUNAGCUCAGCA-GGD--AGAGCAACCGGCUGUU6ACCGGUUU-------------------7UCACAGGUPCGAGCCCUGUAAAAGCCGCCA
=================================================
```

#### Cys

```
>tdbR00000020|Escherichia_coli|562|Cys|GCA
-GGCGCGU4AACAAAGC--GGD--DAUGUAGCGGAPUGCA*APCCGUCU-------------------A-GUCCGGTPCGACUCCGGAACGCGCCUCCA

>tdbR00000019|Mycoplasma_capricolum|2095|Cys|GCA
-GGCAACANGGCCAAGC--GGCD-A"GGCAUGGGUCUGCA=CACCCUGA-------------------U-CAUCGGUPCGAAUCCGAUUGUUGCCUCCA
=================================================
```

63-72

### Ile

```
>tdbR00000156|Bacillus_subtilis|1423|Ile|}AU
-GGACCUUUAGCUCAGUD-GGDD-AGAGCAGACGGCU}AU=ACCGUCCG-------------------7UCGUAGGTPCGAGUCCUACAAGGUCCACCA
>tdbR00000160|Escherichia_coli|562|Ile|}AU
-GGCCCCU4AGCUCAGU--#GDD-AGAGCAGGCGACU}AU6APCGCUUG-------------------7XCGCUGGTPCAAGUCCAGCAGGGGCCACCA
>tdbR00000157|Thermus_thermophilus|274|Ile|GAU
-GGGCGAUUAGCUCAGCU-#GUD-AGAGCGCACGCCUGAU6AGCGUGAG-------------------7UCGGUGGFPCA"GUCCACCAUCGCCCACCA

>tdbR00000153|Mycoplasma_capricolum|2095|Ile|}AU
-GGACCUUNAGCUCAGUD-GGDD-AGAGCAUCCGGCU}AU=ACCGGACG-------------------7UCAUUGGUPCAAGUCCAAUAAGGUCCACCA
>tdbR00000155|Mycoplasma_mycoides|2102|Ile|GAU
-CGGAAUA4AGCUCAGCD-GGDD-AGAGCAUUCCGCUGAU6ACGGAGAG-------------------7UCGUUGGUPCAAGUCCAAUUAUUCCGACCA
=================================================
```

### Phe

```
>tdbR00000065|Bacillus_subtilis|1423|Phe|#AA
-GGCUCGGUAGCUCAGUD-GGD--AGAGCAACGGACU#AA*APCCGUGU-------------------7UCGGCGGTPCGAUUCCGUCCCGAGCCACCA
>tdbR00000067|Escherichia_coli|562|Phe|GAA
-GCCCGGA4AGCUCAGDC-GGD--AGAGCAGGGGAPUGAA*APCCCCGU-------------------7XCCUUGGTPCGAUUCCGAGUCCGGGCACCA
>tdbR00000068|Rhodospirillum_rubrum|1085|Phe|GAA
-GCCCGGGUAGCUCAGCD-GGD--AGAGCACGUGACUGAA*APCACGGU-------------------7UCGGUGGTPCGACUCCGCCCCCGGGCACCA
>tdbR00000069|Synechococcus_sp._PCC_7002|32049|Phe|GAA
-GCCAGGAUAGCNCAGUD-#GD--AGAGCAGAGGACUGAA*APCCUCGU-------------------7UCGGCGGTPCAAUUCCGCCUCCCGGCACCA
>tdbR00000066|Thermus_thermophilus|274|Phe|GAA
-GCCGALG4AGCUCAGUU-#GD--AGAGCAUGCGACUGAA+APCGCAGU-------------------7UCGGCGGTPCGAUUCCGCUCCUCGGCACCA

>tdbR00000064|Mycoplasma_capricolum|2095|Phe|GAA
-GGUCGUGUAGCUCAGUC-GGD--AGAGCAGCAGACUGAAKCPCUGCGU-------------------7UCGGCGGUPCAAUUCCGUCCACGACCACCA
=================================================
```

### Thr

```
>tdbR00000436|Bacillus_subtilis|1423|Thr|5GU
-GCCGGUGUAGCUCAAUD-GGD--AGAGCAACUGACU5GU6APCAGUAG-------------------7UUGGGGGTPCAAGUCCUCUUGCCGGCACCA
>tdbR00000437|Escherichia_coli|562|Thr|GGU
-GCUGAUAUAGCUCAGDD-GGD--AGAGCGCACCCUUGGUEAGGGUGAG-------------------7UCGGCAGTPCGAAUCUGCCUAUCAGCACCA
>tdbR00000438|Escherichia_coli|562|Thr|GGU
-GCUGAUAUGGCUCAGDD-GGD--AGAGCGCACCCUUGGUEAGGGUGAG-------------------7-UCCCAGTPCGACUCUGGGUAUCAGCACCA

>tdbR00000433|Mycoplasma_capricolum|2095|Thr|AGU
-GCUGACUNAGCUCAGUD-GGD--AGAGCAAUUGACUAGU6APCAAUAG-------------------7UCGAAGGUPCAAAUCCUUUAGUCAGCACCA
>tdbR00000434|Mycoplasma_capricolum|2095|Thr|UGU
-GCUGACUNAGCUCAGCA-GGC--AGAGCAACUGACUUGU6APCAGUAG-------------------7UCGUAGGUPCGAUUCCUAUAGUCAGCACCA
=================================================
```

### Trp

```
>tdbR00000483|Bacillus_subtilis|1423|Trp|CCA
-AGGGGCAUAGUUUAAC--GGD--AGAACAGAGGPCUCCA+AACCUCCG-------------------G-UGUGGGTPCGAUUCCUACUGCCCCUGCCA
>tdbR00000484|Escherichia_coli|562|Trp|CCA
-AGGGGCG4AGUUCAADD-GGD--AGAGCACCGGUBUCCA*AACCGGGU-------------------7UUGGGAGTPCGAGUCUCUCCGCCCCUGCCA
>tdbR00000612|Sinorhizobium_meliloti|382|Trp|CCA
-AGGGGUAUAGCUCAGUU-GGU--AGAGCGGCGGUCUCCAAAACCGCAG-------------------GUCGGGGGUUCGAGCCCCUCUGCCCCUGCCA
>tdbR00000482|Spiroplasma_citri|2133|Trp|CCA
-AGGGGUGUAGUUUAAU--GGU--AGAACAGCGGUCUCCAHCACCGUAC-------------------GUUGUGGGUPCAAGUCCUGUCACCCCUGCCA
>tdbR00000481|Spiroplasma_citri|2133|Trp|NCA
-AGGGGUAUAGUUCAAUC-GGU--AGAACACCGGACUNCAHAPCCGGGU-------------------7UUGUGGGUPCAAGUCCUGCUACCCCUGCCA

>tdbR00000480|Mycoplasma_capricolum|2095|Trp|)CA
-AGGGGCAUAGUUCAGUA-GGD--AGAACAUCGGUCU)CA=AACCGAGU-------------------7UCACGAGUPCGAGUCUUGUUGCCCCUGCCA
>tdbR00000479|Mycoplasma_capricolum|2095|Trp|BCA
-AGGAGAGUAGUUCAAU--GGD--AGAACGUCGGUCUBCA=AACCGAGC-------------------7UUGAGGGUPCGAUUCCUUUCUCUCCUGCCA
=================================================
```

**DataSet S2: Archaea- occurrence of coding triplets in the pre-3'end string.** All the unique pre-3'end strings (nucleotides 63-72) of archaeal elongator-tRNAs (not genes) sequences from the tRNAdb[1,2], http://trna.bioinf.uni-leipzig.de, belonging to the 20 amino acids, grouped according to the conservation type determined from the bacterial gene-sequences. The cognate C/ACs conserved in bacteria are boxed and cognate C/ACs not conserved in bacteria are marked by an ellipsoid.

**Archaea- Conserving (in bacteria) amino acids:**

63-72

### Ala

```
>tdbR00000001|Halobacterium_salinarum|2242|Ala|CGC
-GGGCUCGUAGAUCAGC--GGU--AGAUCRCUUCCUUCGCAAGGAAGAG-------------GCC?UGGG]PBOAAUCCCAGCGAGUCCACCA
>tdbR00000002|Haloferax_volcanii|2246|Ala|CGC
-GGGCUCGUAGAUCAGU--GGC--AGAUCRCUUCCUUCGCAAGGAAGAG-------------GC??GGGG]PBOAAUCCCCGCGAGUCCACCA
>tdbR00000003|Haloferax_volcanii|2246|Ala|GGC
-GGGCUCGUAGAUCAGG--GGU--AGAUCACUCCCUUGGCAUGGGAGAG-------------GC??CGGG]PBOAAUCCCGGCCAGUCCACCA
>tdbR00000004|Haloferax_volcanii|2246|Ala|UGC
-GGGCCCAUAGCUCAGU--GGU--AGAGULCCUCCUUUGCAAGGAGGAU-------------GC??AGGG]PBGAAUCCCUGUGGGUCCACCA
==================================================
```

### Arg

```
>tdbR00000589|Aeropyrum_pernix|56636|Arg|UCU
-GGGCCCGUAGCUCAGCCAGGAC-AGAGCGCCGGCCUUCUAAGCCGGUG-------------CUGCCGGGUUCAAAUCCCGGCGGGCCCGCCA
>tdbR00000349|Haloferax_volcanii|2246|Arg|CCG
-GGGCCCGUAGCUCA(U--GGAC-AGAGURCUUGGUUCCGKACCAAGAU-------------GC?GCGGG]PBOAAUCCCGUCGGGUCCGCCA
>tdbR00000350|Haloferax_volcanii|2246|Arg|GCG
-GUCCUGAUARGGPAGU--GGACUAUCCUCCUGGCUUGCGKAGCCAGGG-------------A-C?GGAG]PBOAAUCUCCGUCAGGACGCCA
>tdbR00000351|Haloferax_volcanii|2246|Arg|NCG
-GGGCGCUUAGCUCA(UCUGGAC-AGAGULCUUGGCUNCGKACCAAGUU-------------GC?ACGGG]PBOAAUCCUGUAGCGCCCACCA
>tdbR00000348|Halobacterium_salinarum|2242|Arg|GCG
-GUCCGGAUARGGPAGU--GGACUAUCCUCUUGGCUUGCGKAGCCAGGG-------------A-CCGG?G]PBOAAUCGCCGUCCGGACGCCA
==================================================
```

### Asp

```
>tdbR00000588|Aeropyrum_pernix|56636|Asp|GUC
-GCCGCGGUAGUAUAGCCUGGACUAGUAUGCGGGCCUGUCAAGCCCGUG-------------A-CCCGGGUUCAAAUCCCGGCCGCGGCGCCA
>tdbR00000023|Haloferax_volcanii|2246|Asp|GUC
-GCCCGGGUGRUGPAGU--GGCCCAUCAUACGACCCUGUCACGGUCGUG-------------A-CGCGGG]PBOAAUCCCGCCUCGGGCGCCA
==================================================
```

### Gly

```
>tdbR00000105|Halobacterium_salinarum|2242|Gly|GCC
-GCGCUGGUALUGPAGU--GGU--AUCACGUGACCUUGCCAUGGUCACA-------------A-??UGGG]PBOAAUCCCAGCCAGCGCACCA
>tdbR00000106|Haloferax_volcanii|2246|Gly|CCC
-GCGCCGAUGLUCCAGU--GGU--AGGACACGAGCUUCCCAAGCUCGGA-------------G-C?CGGG]PBOAUUCCCGGUCGGCGCACCA
>tdbR00000107|Haloferax_volcanii|2246|Gly|GCC
-GCGUCGGUALUGPAGU--GGU--AUCACGUGACCUUGCCAUGGUCACA-------------A-C?UGGG]PBOAAUCCCAGCCGACGCACCA
>tdbR00000109|Haloferax_volcanii|2246|Gly|NCC
-GCACCGGUGLUCUAAU--GGU--AAGACAUUGGCCUNCCAAGCCAAUU-------------A-U?UGGG]PBGAUUCCCAGCCGGUGCACCA
>tdbR00000110|Methanobacterium_thermaggregans|83982|Gly|GCC
-GCGGCGUUAGUCCAXCU-GGU-UAAGACACUGGCCUGCCACGCCAGCG-------------U-CCCGGGPPBOAAUCCCGGACGCCGCACCA
==================================================
```

### His

```
>tdbR00000138|Haloferax_volcanii|2246|His|GUG
GUCCGGGUUGRGGPAGU--GGACUAUCCUUCAGCCUUGUGKAGGCUGAG-------------A-CGCGGGPPBAAUUCUCGCACCUGGACCCA
>tdbR00000137|Halobacterium_salinarum|2242|His|GUG
GUCCGGGCU.RGGPAGU--GGACUAUCCUUCAGCCUUGUGKAGGCUGAG-------------A-CGCGGG]PBGAUUCUCGCGCCUGGACCCA
==================================================
```

### Leu

```
>tdbR00000215|Haloferax_volcanii|2246|Leu|5AG
-GCGCGGGUAGCCAA(U--GGCCAAAGGCRCAGCGCU5AGKACGCUGU-GGU-GUAG--ACCUU?GCAGG]PBGAACCCUGUCCCGCGCACCA
>tdbR00000211|Haloferax_volcanii|2246|Leu|CAA
-GCGAGGGUAGCUAA(UCAGGAA-AAAGCRGCGGACUCAAKAPCCGCU-CCC-GUAG--GGGUC?GUGGG]PBOAAUCCCUCCCCUCGCACCA
>tdbR00000212|Haloferax_volcanii|2246|Leu|CAG
-GCAGGGAUAGCCAA(UCUGGCCAACGGCRCAGCGUUCAGKGCGCUGU-CUC-AUAG--GAGUC?GCAGG]PBOAAUCCUGCUCCCUGCACCA
>tdbR00000213|Haloferax_volcanii|2246|Leu|GAG
-GCGUGGGUAGCCAA(CCAGGCCAACGGCRCAGCGUUGAGKG?GCUGU-CCU-GUAG--AGGUC?GCCGG]PBOAAUCCGGUCCCACGCACCA
>tdbR00000214|Haloferax_volcanii|2246|Leu|UAA
-GCGGGGGUGGCUGA(CCAGGCCAAAAGCLGCGGACUUAAKAPCCGCU-CCC-GUAG--GGGUUCGCGAG]PBGAAUCUCGUCCCCCGCACCA
```

**Pro**

63-72

```
>tdbR00000579|Aeropyrum_pernix|56636|Pro|CGG
-GGGCCCGUCGUCUAGCCUGGCU-AAGAUGCGGGGUACGGACCCCGUG-------------GUCCGGGGUUCAAAUCCCCGCGGGCCCACCA
>tdbR00000584|Aeropyrum_pernix|56636|Pro|GGG
-GGGGCCGUCGUCUAGCCUGGCU-AGGAUGCCAGCCUGGGGCGCUGGUG-------------GUCCCGGGUUCAAAUCCCGGCGGCCCACCA
>tdbR00000312|Haloferax_volcanii|2246|Pro|GGG
-GGGACCGUGRGGPAGU--GGU--AUCCUCUGCCGAUGGGKUCGGUAGG-------------A-C?UGAG]PBGACUCUCAGCGGUCCCACCA
>tdbR00000311|Haloferax_volcanii|2246|Pro|MGG
-GGGCCGGUGRGGPA(CUUGGU--AUCCUUCGGCCUUMGGKPGGCCGUA-------------A-??UCAG]PBGAAUCUGAGCCGGCCCACCA
>tdbR00000313|Haloferax_volcanii|2246|Pro|UGG
-GGGACCGUGRGUPA(CCUGGU--AUACUUCGGGCCUUGGKUGCCCGUG-------------A-??CCGG]PBOAAUCCGGGCGGUCCCACCA
=================================================
```

**Ser**

```
>tdbR00000381|Haloferax_volcanii|2246|Ser|GCU
-GUUGCGGUAGCCAA(CCUGGCCCAAGGCRCUGGGUUGCU6ACUCAGU-GGC-GUCAA-GCCC-??GGGG]PBGAAUCCCCGCCGCAACGCCA
>tdbR00000383|Haloferax_volcanii|2246|Ser|GGA
-GCCAGGAUGGCCGA(C--GGU--AAGGCRCACGCCUGGAAAGCGUGU-UCCCUCU--GGGAU-?GGGGG]PBOAAUCCCUCUCCUGGCGCCA
>tdbR00000382|Haloferax_volcanii|2246|Ser|MGA
-GCCGAGGUAGCCPA(CCCGGCC-AAGGCRGUAGAUUMGAAAPCUACU-GUCCAUUC-GGACA-?GUGAG]PBOAAUCUCACCCUCGGCGCCA
>tdbR00000380|Halobacterium_salinarum|2242|Ser|MGA
-GCCGAGGUAGCAPA(CUUGGCC-AAUGCRGUUGCUUMGAKAGCAACG-UUCCACAC-GGACU-?AGGAG]PBOAAUCUCCUCCUCGGCGCCA
=================================================
```

**Val**

```
>tdbR00000446|Halobacterium_salinarum|2242|Val|CAC
-GGGUCGGUGGUCPAGUCCGGUU-AUGACGGCUCCUUCACACGGAGCAG-------------GUCGGCGG]P#OACUCCGCCCCGACCCACCA
>tdbR00000447|Halobacterium_salinarum|2242|Val|CAC
-GGGUUGGUKGUCPAGUCAGGCU-AUGACACCUCCUUCACAUGGAGGAG-------------GUCGG?GG]PBOAAUCCGCCCCAACCCACCA
=================================================
```

**Archaea- Other amino acids (according to bacteria)**

**Gln**

```
>tdbR00000329|Halobacterium_salinarum|2242|Gln|CUG
-AGUCCCGU.RGGPAGU--GGCCAAUCCUGAAGCCUUCUGKGGGCUUCG-------------A-CGGAAGPPBGAAUCUUCCCGGGACUACCA
>tdbR00000330|Haloferax_volcanii|2246|Gln|MUG
-AGUCCCAUGRGGPAGU--GGCCAAUCCUGUUGCCUUMUGKGGGCAACG-------------A-CCCAGGPPBGAAUCCUGGUGGGACUACCA
>tdbR00000617|Nanoarchaeum_equitans|160232|Gln|UUG
-AGCCCCGUGGUGUAGC--GGCCUAGCAUGCGGGGUUUUGGUCCCCGUG-------------A-CCCCGGUUCGAAUCCGGGCGGGGCUACCA
=================================================
```

**Glu**

```
>tdbR00000613|Nanoarchaeum_equitans|160232|Glu|CUC
-GCCCCCGUGGUGUAGCCAGGUCUAGCAUACGGGGUUCUCGUCCCCGUG-------------A-CCCGGGUUCAAAUCCCGGCGGGGGCACCA
>tdbR00000042|Haloferax_volcanii|2246|Glu|MUC
-GCUCUGUUGRUGPAGUCCGGCCAAUCAUAUCACCCUMUCACGGUGAUG-------------A-C?AGGG]PBGAAUCCCUGACGGAGCACCA
>tdbR00000043|Haloferax_volcanii|2246|Glu|NUC
-GCUCGGUUGRUGPAGUCCGGCCAAUCAUCUUGGCCUNUCKAGCCGAGG-------------A-C?AGGG]PBOAAUCCCUGACCGAGCACCA
=================================================
```

**Lys**

```
>tdbR00000176|Haloferax_volcanii|2246|Lys|MUU
-GGGCCGGUAGCUCA(UUAGGC--AGAGCRUCUGABUMUU6APCAGACG-------------GU?GCGPG]PBOAAUCGCGUCCGGCCCACCA
>tdbR00000175|Haloferax_volcanii|2246|Lys|NUU
-GGGCUGGUAGCUCA(UUAGGC--AGAGCRUCUGGBUNUU6ACCAGACG-------------GU?GGGGG]PBOAGUCCCUCCCAGCCCGCCA
>tdbR00000581|Aeropyrum_pernix|56636|Lys|CUU
-GGGCCCGUAGCUCAGCCUGGU--AGAGCGGCGGGCUCUUAACCCGCGAGG--GAGG---AAGUCCCGGGUUCAAAUCCCGGCGGGCCCGCCA
>tdbR00000616|Nanoarchaeum_equitans|160232|Lys|CUU
-GGGCCGGUGGCUCAGCCUGGUU-AGAGCGGCGGGCUCUUAACCCGCAG-------------GUCCGGGGUUCGAAUCCCCGCCGGCCCGCCA
=================================================
```

**Met**

```
>tdbR00000586|Aeropyrum_pernix|56636|Met|CAU
-GGGCCCGUAGCUCAGCCAGGU--AGAGCGCCCGGCUCAUAACCGGGUG-------------GUCGGGGGUUCAAAUCCCCCCGGGCCCACCA
>tdbR00000587|Aeropyrum_pernix|56636|Met|CAU
-GCCGCCGUAGCUCAGCCUGGC--AGAGCGCCGGACUCAUAAUCCGGAG-------------GUCCCGGGUUCGAAUCCCGGCGGCGGCACCA
>tdbR00000271|Haloferax_volcanii|2246|Met|BAU
-GCCCGGGUGGCUPA(CU-GGAC-APAGCGCCGCACUBAU6APGCGGAG-------------AU?GUGGG]PBGGAGCCCACCCCGGGCACCA
>tdbR00000272|Thermoplasma_acidophilum|2303|Met|CAU
-GCCGGGG4GGCUCA(CU-GGA--GGAGCRCCGGABUCAU6AUCCGGAG-------------GUCUCGGGPPBGAUCCCCGAUCCCGGCACCA
=================================================
```

**Tyr**

```
>tdbR00000541|Haloferax_volcanii|2246|Tyr|GUA
-CCGCUCUUAGCUCA(CCUGGC--AGAGCAGCCGABUGUAKAPCGGCUU-------------GU?CCCCG]PBOAAUCGGGGAGAGCGGACCA
>tdbR00000582|Aeropyrum_pernix|56636|Tyr|GUA
-CCCGCCGUAGCUCAGC--GGC--AGAGCGCCCGGCUGUACACCGGGUG-------------GUCCGGGGUUCGAAUCCCCGCGGCGGGACCA
================================================
```

63-72

### Asn

```
>tdbR00000291|Halobacterium_salinarum|2242|Asn|GUU
GCCGCCAUAGCUCAGUU-GGU--AGAGCACGUGGUUGUU6CCCACGUU-------------GU?CCAGG]PBGGAC CCUGGUGGCGGCGAAC
>tdbR00000292|Haloferax_volcanii|2246|Asn|GUU
-GCCGCCGUAGCUCA(UU-GGU--AGAGCACCUCGCUGUU6ACGAGGUU-------------GU??CAGG]PBGAGUCCUGGCGGUGGCGCCA
>tdbR00000293|Methanobacterium_thermaggregans|83982|Asn|;UU
-GCGCCGGUGGCUCAXCCUGGUU-AGAGCUCACGGCU;UU6ACCGUGAG-------------GCCGCGGGPPBOAAUCCCGCCCGGCGCACCA
==================================================
```

### Cys

```
>tdbR00000018|Haloferax_volcanii|2246|Cys|GCA
-GCCAAGGUGGCAGA(UUCGGCCCAACGCAUCCGCCUGCAKAGCGGAAC-------------C_?GCCGG]PBOAAUCCGGCCCUUGGCUCCA
>tdbR00000583|Aeropyrum_pernix|56636|Cys|GCA
-GCCGGGGUGGCCGAGC--GGUCUAAGGCGGCGGGCUGCAGACCCGUUA----G--------UUCCCGGGUUCGAAUCCCGGCCCCGGCUCCA
==================================================
```

### Ile

```
>tdbR00000151|Haloferax_volcanii|2246|Ile|GAU
-GGGCCAAUAGCUCAGUCAGGUU--GAGCRCPCGGCUGAU6AC?GGGAG-------------GCC?GCGG]PBOAAUCCGCGUUGGCCCACCA
>tdbR00000152|Haloferax_volcanii|2246|Ile|NAU
-GGGCCCCUAGCUCA(UCUGGUC-AGAGCRCUCGGCUNAU6ACCGGGUG-------------GU?AUGGG]PBGAACCCCAUGGGGCCCACCA
==================================================
```

### Phe

```
>tdbR00000063|Haloferax_volcanii|2246|Phe|GAA
-GCCGCCUUAGCUCA(ACUGGG--AGAGCACUCGACUGAAKAPCGAGCU-------------GU?CCCGG]PBOAAUCCGGGAGGCGGCACCA
==================================================
```

### Thr

```
>tdbR00000431|Haloferax_volcanii|2246|Thr|CGU
-GCCGGUGUAGCUCA(UU-GGC--AGAGCRAUUCCUUCGU6AGGAAUAG-------------GC?GAGGG]PBOAAUCCCUCCACCGGCUCCA
>tdbR00000585|Aeropyrum_pernix|56636|Thr|CGU
-GCCGCCGUAGCUCAGC--GGU--AGAGCGCCGGCCUCGUAAGCCGGUG-------------GUCGCGGGUUCGAAUCCCGCCGGCGGCUCCA
>tdbR00000430|Halobacterium_salinarum|2242|Thr|GGU
-GCCUGGGUAGCUPAGC--GGU--AAAGCRCGUCCUUGGU6AGGACGAG-------------ACC??GGA]PBOAAUUCCGGCCUAGGCUCCA
==================================================
```

### Trp

```
>tdbR00000478|Haloferax_volcanii|2246|Trp|BCA
-GGGGCUGUGGCCAA(CCCGGC--AUGGCRACUGABUBCAKAJCAGUCG-------------AU?GGGGG]PBOAAUCCCUCCGGCCCCACCA
>tdbR00000615|Nanoarchaeum_equitans|160232|Trp|CCA
-GGGGCCGUAGCUCAGCCAGGC--AGAGCGGCGGGCUCCAGACCCGUAG-------------GUCGGGGGUUCGAAUCCCCCCGGCCCCACCA
==================================================
```

**DataSet S3: Eukarya- occurrence of coding triplets in the pre-3'end string.** All the unique pre-3'end strings (nucleotides 63-72) of eukaryotic elongator-tRNAs (not genes) sequences from the tRNAdb[1,2], http://trna.bioinf.uni-leipzig.de, belonging to the 20 amino acids, grouped according to the conservation type determined from the bacterial gene-sequences. The cognate C/ACs conserved in bacteria are boxed and cognate C/ACs not conserved in bacteria are marked by an ellipsoid.

### Eukarya- Conserving (in bacteria) amino acids:

#### Ala

63-72

```
>tdbR00000014|Bombyx_mori|7091|Ala|IGC
-GGGGGCGUALCUCAGAD-GGU--AGAGCRCUCGCJUIGCOP#PGAGAG--------------7UA?CGGGAPCG"UACCCGGCGCCUCCACCA
>tdbR00000016|Homo_sapiens|9606|Ala|IGC
-GGGGGAUUALCUCAAAD-GGD--AGAGCRCUCGCJUIGCOP#CGAGAG--------------7UAGCGGGAPCG"UGCCCGCAUCCUCCACCA
>tdbR00000013|Pichia_jadinii|4903|Ala|IGC
-GGGCGUGUKGCGUAGDD-GGD--AGCGCRPUCGCUUIGCOPGCGAAAG--------------GDCUCCGGTPCG"CUCCGGACUCGUCCACCA
======================================================
```

#### Arg

```
>tdbR00000376|Bos_taurus|9913|Arg|3CP
-GGCUCCGUKGCGCAAD--GGAD-AGCGCAPPGGA'U3CP6AUPCAAAG--------------7DU?CGGGTPCG"GUCCCGGCGGAGUCGCCA
>tdbR00000374|Bos_taurus|9913|Arg|CCG
-GACCCAGUKLCCUAAD--#GAD-AAGGCAPCAGCBUCCGKAGCUGGGG--------------ADUGPGGGTPCG"GUCCCAUCUGGGUCGCCA
>tdbR00000377|Mus_musculus|10090|Arg|ICG
-GGGCCAGUKLCGCAAD--GGAD-AACGCRPCUGABUICGKAPCAGAAG--------------ADU?PAGGPPCG"CUCCUGGCUGGCUCGCCA
>tdbR00000370|Saccharomyces_cerevisiae|4932|Arg|1CU
-GCUCGCGUKLCGUAAD--GGC--AACGCRPCUGACU1CU6APCAGAAG--------------ADUAUGGGTPCG"CCCCCAUCGUGAGUGCCA
>tdbR00000369|Saccharomyces_cerevisiae|4932|Arg|ICG
 -PUCCUCGUKLCCCAAD—GGDC-ACGGCRPCUGGCUICGAACCAGAAG--------------ADU?CAGGTPCA"GUCCUGGCGGGAAGCCA
>tdbR00000373|Triticum_aestivum|4565|Arg|ICG
-GACUCCGUKLCCCAAD--#GAX-AAGGCRCUGGUBUICG/AACCAGAG--------------ADU?UGGGTPCG"UCCCCAGCGGAGUCGCCA
========================================================
```

#### Asp

```
>tdbR00000040|Bos_taurus|9913|Asp|8UC
 -GGUGCCGUALCGPAGD--#GC—A.CG.GACUCUBU8UCAAAGAGUGG--------------7A?.UGAGTPCG"UACUCAACGGCACCGCCA
>tdbR00000036|Euglena_gracilis|3039|Asp|GUC
-UCUUCGGUAGUAPAGD--#GDA-AGUAU;PCCGCCUGUCA<GCGGAAG--------------A-<HCGGGTPCAHUUCCCGGCCGGAGAGCCA
>tdbR00000039|Oryctolagus_cuniculus|9986|Asp|8UC
 -UCCCGUCUAGUAPAGU—GGDG-AGUAUAUCCGCCU8UCACGCGGGAG--------------GA??GGGGTPCG"AUCCCCGGACGGGAGCCA
>tdbR00000037|Rattus_norvegicus|10116|Asp|GUC
 AUCCUCGUUAGUAPAGU—GGDG-AGUAUCCCCGCUCGUCA?GCGGGAG--------------A-??GGGGTPCGAUUCCCCGACGGGGAGCCA
>tdbR00000035|Saccharomyces_cerevisiae|4932|Asp|GUC
 -UCCGUGAUAGUUPAAD—GGDC-AGAAUGGGCGCPUGUCKCGUGCCAG--------------A-U?GGGGTPCAAUUCCCCGUCGCGGAGCCA
====================================================
```

#### Gly

```
>tdbR00000134|Bombyx_mori|7091|Gly|,CC
 -GCGJUGGUKGUGPAAD—GGDC-AGCAUAGPUGCCU,CCAAGCAGUUG--------------A-U??GGGTPCG"UUCCGGCCAACGCACCA
>tdbR00000133|Bombyx_mori|7091|Gly|GCC
  -GCAJCGGUKGUUCAGU—GGD—AGAAUGCUCGCCUGCCA?GCGGGCG--------------G-???GGGTPCG"UUCCCGGCCGAUGCACCA
>tdbR00000135|Homo_sapiens|9606|Gly|CCC
  -GCGBCLCUGGUGPAGU—GGD—AUCAUGCAAGAJUCCCAUZCUUGCG--------------A-C??GGGTPCG"UUCCCGGGGGCGCACCA
>tdbR00000136|Homo_sapiens|9606|Gly|GCC
  -GCAJULGUGGUUCAGU—GGD—AGAAUUCUCGCCUGCCA?GCGGGAG--------------G-???GGGTPCG"UUCCCGGCCAAUGCACCA
>tdbR00000132|Lupinus_luteus|3873|Gly|GCC
  -GCABCAGUKGUCPAGD—GGU—AGAAUAGUACCCUGCCA?GGUACAG--------------A-???GGGUPCG"UUCCCGGCUGGUGCACCA
>tdbR00000130|Saccharomyces_cerevisiae|4932|Gly|.CC
 -GGGCGGUUAGUGPAGD—GGDD-AUCAUCCCACCCU.CCAAGGUGGGG--------------A-CACGGGTPCGAUUCUCGUACCGCUCACCA
>tdbR00000129|Saccharomyces_cerevisiae|4932|Gly|GCC
  -GCGBAAGUKGUUPAGD—GGD—AAAAUCCAACGPUGCCAPCGUUGGG----------------C?CCGGTPCGAUUCCGGGCUUGCGCACCA
>tdbR00000131|Triticum_aestivum|4565|Gly|GCC
  -GCABCAGUKGUCPAGD—GGU—AGAAUAGUACCCUGCCA?GGUACAG--------------A-???GGGUPCG"UUCCCGGCUGGUGCACCA
====================================================
```

#### His

```
>tdbR00000149|Homo_sapiens|9606|His|GUG
 GGCCGUGAUCGUAPAGD—GGDD-AGUACUCUGCGPUGUGGCCGCAGCA--------------A-??UCGGUPCG"AUCCGAGUCACGGCACCA
>tdbR00000146|Lupinus_luteus|3873|His|GUG
 GGUGGCUGUAGUUPAGD—GGDD-AGAACACAACGPUGUGKCCGUUGAA--------------A-C?UGGGUPCG"AUCCCAGCAGCCACACCA
>tdbR00000144|Saccharomyces_cerevisiae|4932|His|GUG
GGCC:UCUUAGUAPAGD--#GDD-AGUACACAACAPUGUGKCPGUUGAA--------------A-C?CUGGTPCGAUUCUAGGAGGUGGCACCA
>tdbR00000145|Saccharomyces_cerevisiae|4932|His|GUG
GGCC:UCUUAGUAPAGD--#GDD-AGUACACAUCGPUGUGKCCGAUGAA--------------A-C?CUGGTPCGAUUCUAGGAGAUGGCACCA
===================================================
```

### Leu

63-72

```
>tdbR00000265|Bos_taurus|9913|Leu|IAG
 -GGUAGCGUGLCMGAGC—GGDCPAAGGCRCUGGAZUIAGKCPCCAGU-CPCP-UC---GGGGG-?GUGGGTPCG”AUCCCACCGCUGCCACCA
>tdbR00000266|Bos_taurus|9913|Leu|°AA
 -GUCAGLAUGLCMGAGU—GGDCPAAGGCLCCAGACU°AAKPPCUGGJ-CPCC-GUAU-GGAGG-?GUGGGTPCG”AUCCCACUUCUGACACCA
>tdbR00000264|Caenorhabditis_elegans|6239|Leu|IAG
 -GGAGAGAUGGCMGAGC—GGDCUAAGGCGCUGGUUUIAGGCACCAGU-CCCU-UC---GGGGG-CGUGGGTUCGAAUCCCACUCUCUUCACCA
>tdbR00000618|Candida_albicans|5476|Leu|CAG
 -GAUACGAUGGCMGAGD--#GDD-AAGGCRAAGGAUGCAGKPPCCUU—UGGGCAUU-GCCCG--?GCAGGTPCG”ACCCUGCUCGUGUCGCCA
>tdbR00000253|Candida_cylindracea|44322|Leu|BAA
  -GGCCGUUUGLCMGAGD--#GDCDAAGGCRUCUGABUBAAKAPCAGAJ-CUC—GUAA—GAGG-?GUGUGTPCG”ACCACACAGCGGUCACCA
>tdbR00000254|Candida_cylindracea|44322|Leu|IAG
   -GGUUCUCUGGCMGAGD—GGDCDAAGGCRCAUGGPUIAGKPCCAUGU-CUC—UUCG—GAGG-?GCGAGTPCG”ACCUCGCGGGAAUCACCA
>tdbR00000255|Candida_cylindracea|44322|Leu|IAG
   -GGCUCUCUGGCMGAGD—GGDCDAAGGCRCUAGGGUIAGKPCCUAGU-CUC—UUCG—GAGG-?GCGAGTPCG”ACCUCGCGGGAGUCACCA
>tdbR00000605|Lupinus_luteus|3873|Leu|CAG
 -GUCAGGAUGLCMGAG-D-GGDCXAAGGCRCCAGUJUCAGKPACUGGJCPCG—AAC---CGGG-?AUGGGTPCG”AUCCCAUUCUUGACACCA
>tdbR00000606|Lupinus_luteus|3873|Leu|IAG
-GUCAGGAUGLCMGAG-D-GGDCXAAGGCRCCAGAZUIAGKPUCUGGJCPCGA-A----UCGGG-?GUGGGTPCA”AUCCCACUCUUGACACCA
>tdbR00000259|Phaseolus_vulgaris|3885|Leu|.AA
 -GCUGGUUUGGCMGAGA—GGDD-AAGGCRGAAGACU.AAKAPCUUCJ-GCAG-UCAA-CUGCG-?AUGGGTPCG”ACCCCAUAGCCAGCACCA
>tdbR00000258|Phaseolus_vulgaris|3885|Leu|.AG
  -GAUAGUUUGLCMGAGD—GGDCXAAGGCRCCAGAPU.AGKCPCUGGJ-CCG—AAA---.GGG-?GUGGGUPCA”AUCCCACAGCUGUCACCA
>tdbR00000252|Pichia_jadinii|4903|Leu|BAA
  -GGAUCUUUGLCMGAGC--#GDDUAAGGCRCUCGABUBAAKAPCGAGU-AUC—GUAA—GAUG-?AUGAGTPCG”AUCUCAUAGGAUCCACCA
>tdbR00000249|Saccharomyces_cerevisiae|4932|Leu|CAA
  -GGUUGUUUGLCMGAGC--#GDCDAAGGCRCCUGAPU?AAKCPCAGGU-AUC—GUAA—GAUG-?AAGAGTPCGAAUCUCUUAGCAACCACCA
>tdbR00000251|Saccharomyces_cerevisiae|4932|Leu|UAA
-GGAGGGUUGLCMGAGD--#GDCDAAGGCRGCAGABUUAAKAPCUG-UUGGAC-GGUU-GUCC-G?GCGAGTPCG”ACCUCGCAUCCUUCACCA
>tdbR00000250|Saccharomyces_cerevisiae|4932|Leu|UAG
  -GGGAGUUUGLCMGAGD--#GDDDAAGGCRPCAGAPUUAGKCPCUGAU-AUC—UUCG—GAUG-?AAGGGTPCG”AUCCCUUAGCUCUCACCA
==================================================
```

### Pro

```
>tdbR00000326|Lupinus_luteus|3873|Pro|IGG
  -GGGACAUUKGUCPAGD—GGD—AUGAUUCUCGCUUIGGKPGCGAGAG--------------7D?CCGAGTPCG”UUCUCGGAUGUCCCCCCA
>tdbR00000325|Pichia_jadinii|4903|Pro|&GG
  -GGCBGCGUKGUCPAGD—GGD—AUGAUACUCGCUU&GGKPGPGAGUG--------------7D??AGGGTPCA”UUCCCUGCUCGGCCCCCA
>tdbR00000323|Saccharomyces_cerevisiae|4932|Pro|NGG
  -GGGBGUGUKGUCPAGD—GGD—AUGAUUCUCGCPUNGGKPGCGAGAG--------------7CCCUGGGTPCA”UUCCCAGCUCGCCCCCCA
===================================================
```

### Ser

```
>tdbR00000410|Candida_cylindracea|44322|Ser|&GA
 -GGUGCAAUGGCMGAGD--#GDD-AAGGCRACGGAPU&GA+CPCCGUJ-GGGA-UUC—UCCCU-?GCAGGTPCG”AUCCUGUUUGCAUCGCCA
>tdbR00000409|Candida_cylindracea|44322|Ser|CAG
-GACACCGUGLCMCAGD--#GDD-ACGGCRPUGGACGCAGAAPCCAUU-GAGC-UU---GCUCG-?GCGGGTPCG”AUCCCGUCGGUGUCUCCA
>tdbR00000412|Candida_cylindracea|44322|Ser|CGA
-GGCAACUUGGCMGAGD--#GDD-AAGGCRUGACA’UCGA+APGUCAJ-GGGU-UU---ACCCG-?GCAGGTPCG”AUCCUGCAGUUGUCGCCA
>tdbR00000413|Candida_cylindracea|44322|Ser|GCU
 -GUUAAUGUGGCCMGAGD-GGDD-AAGGCRCUUCC’UGCU+AGGAAGJ-GGGC-UCU—GCCCG-=GCAGGTPCG”AUCCUGUCAUUAACGCCA
>tdbR00000411|Candida_cylindracea|44322|Ser|IGA
 -GGCAUCUUGGCMGAGD--#GDD-AAGGCRAAAGAPUIGA+APCUUUJ-GGGC-UCU—GCCCG-?GCAGGTPCG”GUCCUGCAGAUGUCGCCA
>tdbR00000426|Drosophila_melanogaster|7227|Ser|CGA
 -GCA#UCGUGGCMGAGD--#GDD-AAGGCRUCUGA’UCGA+APCAGAJ-UCCC-U’U—GGGAG-?GUAGGTPCG”AUCCUACCGGCUGCGCCA
>tdbR00000424|Drosophila_melanogaster|7227|Ser|GCU
 -GACGAGGUGGCMGAGD--#GDD-AAGGCRPPGGA’UGCUEAPCCAAJ-GUGC-U’U—GCACG-?GUGGGTPCG”AUCCCAUCCUCGUCGCCA
>tdbR00000425|Drosophila_melanogaster|7227|Ser|IGA
 -GCA#UCGUGGCMGAGC--#GDD-AAGGCRUCUGA’UIGA+APCAGAJ-UCCC-U’U—GGGAG-?GUAGGTPCG”AUCCUACCGACUGCGCCA
>tdbR00000427|Gallus_gallus|9031|Ser|UGA
-AAGAAAGA”LCAUUAAGUGGDUUGAUGCLGPUGG’UUGAHACCAACAU--------------G-UGAGGGTPCG”UUCCUUCCUUUCUUGCCA
>tdbR00000428|Homo_sapiens|9606|Ser|UGA
 -GUAGUCGUGGCMGAGD--#GDD-AAGGCGAPGGACUUGAAAPCCAUJ-GGGG-U’U—CCCCG-?GCAGGTPCG”AUCCUGCCGACUACGCCA
>tdbR00000414|Lupinus_luteus|3873|Ser|CGA
 -GUCAGUAUGGCMGAGD--#GDD-AAGGCRACAGA’UCGA`APCUGUJ-GAGAUUU—GCCUGG-?AGCGGTPCG”AUCCGCUUACUGACGCCA
>tdbR00000415|Nicotiana_rustica|4093|Ser|&GA
 -GUCGAUAUGUCCGAGD--#GDD-AAGGARACAGA’U&GA`APCUGUJ-GGGC-UUC—GCCCG-CGCAGGTPCG”ACCCUGCUGUCGACGCCA
>tdbR00000417|Nicotiana_rustica|4093|Ser|GCU
 -GUCGCCUUGGCCGAGD--#GDD-AAGGCRPGUGC’UGCU6AGPACAJ-GGGG-UUU—CCCCG-CGAGAGTPCG”AUCUCUCAGGCGACGCCA
>tdbR00000418|Nicotiana_rustica|4093|Ser|IGA
 -GUGGACGUGCCGGAGD--#GDD-AUCGGRCAUGA’UIGA`APCAUGJ-GGGC-UUU—GCCCG-CGCAGGTPCG”AUCCUGCCGUCCACGCCA
>tdbR00000406|Saccharomyces_cerevisiae|4932|Ser|NGA
 -GGCACUAUGGCMGAGD--#GDD-AAGGCRACAGA<UNGA+APCUGUJ-GGGC-UCU—GCCCG-?GCUGGTPCAAAUCCUGCUGGUGUCGCCA
=======================================================
```

### Val

63-72

```
>tdbR00000473|Drosophila_melanogaster|7227|Val|CAC
 -GUUUCCGUAGUGPAGC—GGDX-AUCACGPGUGCUUCACACGCACAAG--------------7DCCCCGGTPCG”ACCCGGGCGGGAACACCA
>tdbR00000472|Drosophila_melanogaster|7227|Val|IAC
 -GUUJCCGUKGUGPAGC—GGDX-AUCACAPCUGCBUIACA?GCAGAAG--------------7CC<CCGGTPCG”UCCCGGGCGGAAACACCA
>tdbR00000474|Drosophila_melanogaster|7227|Val|NAC
 -GGUJCCAUAGUGPAGC—GGDN-AUCAC;PCUGCPUNACACGCAGAAG--------------7D?UCCGGTPCG”UCCCGGAUGGAACCACCA
>tdbR00000468|Lupinus_luteus|3873|Val|IAC
 -GGUUUCGUKGUGPAGD—GGDD-AUCACRPCAGUCUIACACACUGAAG--------------7DCUCCGGTPCG”GUCCGGGCGAAGCCACCA
>tdbR00000467|Pichia_jadinii|4903|Val|IAC
-GGUUUCGUKGUCPAGDD-GGDC-AUGGCAPCUGCPUIACACGCAGAAC--------------7DC?CCAGTPCG”UCCUGGGCGAAAUCACCA
>tdbR00000466|Saccharomyces_cerevisiae|4932|Val|&AC
 -GGUCCAAUGLUCCAGD—GGDDCAAGACRPCGCCPU&ACACGGCGAAG--------------ADC?CGAGTPCG”ACCUCGGUUGGAUCACCA
>tdbR00000465|Saccharomyces_cerevisiae|4932|Val|CAC
 -GUUCCAAUALUGPAGC—GGCD-AUCACLPPGCCPUCACACGGCAAAG--------------7DC?CGAGTPCG”UCCUCGGUUGGAACACCA
```

### Eukarya-other amino acids (according to bacteria)

#### Gln

```
>tdbR00000342|Bos_taurus|9913|Gln|CUG
-GGUUCCAUKGUGPAAD--#GDD-AGCACUCPGGABUCUGAAPCCAGCG--------------A-U?CGAGPPCA”AUCUCGGUGGAACCUCCA
>tdbR00000345|Homo_sapiens|9606|Gln|NUG
-GGCCCCAUKGUGPAAU--#GDD-AGCACUCPGGABUNUGAAPCCAGCG--------------A-U??GAGPPCA”AUCUCGGUGGGACCUCCA
>tdbR00000604|Lupinus_luteus|3873|Gln|UUA
     -..CGUCUUAL…AGD—GG..-A…CR.C.UGBUUUAEACACG.UG--------------7D?GUGGGTPCG”AUCCCACAGACGAGACCA
>tdbR00000340|Nicotiana_rustica|4093|Gln|JUG
-GGUUCUAUKGUGPAGD--#GDD-AGCACUCPGGABUJUG/APCCAGCG---------------ACCUGGGTPCG”CUCCCGGUAGGACCUCCA
>tdbR00000337|Tetrahymena_thermophila|5911|Gln|CUA
 -GGUUCUAUALUAPAGC—GCDD-AGUACUGGGGA<UCUA6APCCCUUG--------------A-C?UGGGUPCG”AUCCCAGUAGGACCUCCA
>tdbR00000338|Tetrahymena_thermophila|5911|Gln|JUA
 -GGUUCCAUALUAPAGD—GGDD-AGUACUGGGGABUJUA6APCCCUUG--------------A-C?UGGGUPCG”AUCCCAGUGGGACCUCCA
>tdbR00000339|Tetrahymena_thermophila|5911|Gln|JUG
 -GGUUGUAUKLUGPAGC—GGDD-AGCACCGAGGACUJUGAAPCCUCUG--------------A-C?UGGGUPCG”AUCCCAGUACGACCUCCA
=================================================
```

#### Glu

```
>tdbR00000060|Drosophila_melanogaster|7227|Glu|2UC
 -UCCCAUAUGGUCPAGD—GGCD-AGGAUAUCUGGBU2UCACCCAGAAG--------------G-CCCGGGTPCGAUUCCCGGUAUGGGAACCA
>tdbR00000062|Homo_sapiens|9606|Glu|CUC
 -UCCCUGGUGLUCPAGU—GGDP-AGGAUUCGGCGCUCUCACCGCCGCG--------------G-C??GGG\PCGAUUCCCGGUCAGGGAACCA
>tdbR00000057|Hordeum_vulgare|4513|Glu|CPC
-UCCGUCGUAGUCPAGGD-GGDP-AGGAUACUCGGCUCPCACCCGAGAG--------------A-C?CGGGTPCG”GUCCCGGCGACGGAACCA
>tdbR00000059|Rattus_norvegicus|10116|Glu|3UC
 -UCCCACAUKGUCPAGC—GGDD-AGGAUUCCUGGPU3UCACCCAGGCG--------------G-C??GGG\PCGACUCCCGGUGUGGGAACCA
>tdbR00000054|Saccharomyces_cerevisiae|4932|Glu|3UC
 -UCCGAUAUAGUGPAAC—GGCD-AUCACAPCACGCU3UCACCGUGGAG--------------A-C?GGGGTPCGACUCCCCGUAUCGGAGCCA
>tdbR00000055|Schizosaccharomyces_pombe|4896|Glu|3UC
 -UCCGUUGUKGUCCAAC—GGCD-AGGAUUCGUCGCU3UCACCGACGGG--------------A-G?GGGGTPCGACUCCCCGCAACGGAGCCA
==================================================
```

#### Lys

```
>tdbR00000202|Drosophila_melanogaster|7227|Lys|)UU
 -GCCCGGAUALCUCAGDC-GGD—AGAGCAPPGGACU)UU6APCCAAGG--------------7D?CAGGG\PCA”GUCCCUGUUCGGGCGCCA
>tdbR00000201|Drosophila_melanogaster|7227|Lys|CUU
 -GCCCGGCUALCUCAGDC-GGD—AGAGCAPGAGACUCUU6APCUCAGG--------------7DCGUGGG\PCG”GCCCCACGUUGGGCGCCA
>tdbR00000603|Homo_sapiens|9606|Lys|tUU
 -CACUGUAA”LCU-AAC----U—U-AGCAPPAACCUtUU6AGUUAAAG--------------AUUAAGAGAACCA__CUCUUUACAGUGACCA
>tdbR00000204|Loligo_bleekeri|6617|Lys|CUU
 -UCCCGLCUALCUCAGDC-GGU—AGAGCACGAGANUCUU6APCUCGGG--------------7D?GUGGGJPCG”GCCCCACGUUGGGAGCCA
>tdbR00000192|Saccharomyces_cerevisiae|4932|Lys|CUU
 -GCCUUGUULLCGCAADC-GGD—AGCGCRPAUGACUCUU6APCAUAAG--------------7UUAGGGGTPCG”GCCCCCUACAGGGCUCCA
================================================
```

#### Met

```
>tdbR00000289|Homo_sapiens|9606|Met|BAU
 -GCCUCLUUALCGCAGDA-GGD—AGCGCRPCAGPCUBAU6APCUGAAG--------------7D?GUGAGTPCG”UCCUCACACGGGGCACCA
>tdbR00000286|Lupinus_luteus|3873|Met|BAU
 GGGGUGGUKLCGCAGDD-GGCX-AGCGCRPAGGPCUBAUEAPCCUGAG--------------7D?GAGAGTPCG”GCCUCUCUCACCCCACCA
>tdbR00000284|Saccharomyces_cerevisiae|4932|Met|CAU
 -GCUUCAGUALCUCAGDA-GGA—AGAGCRPCAGPCUCAU6APCUGAAG--------------7D?GAGAGTPCG”ACCUCUCCUGGAGCACCA
>tdbR00000602|Tetrahymena_pyriformis|5908|Met|CAU
  -GCUGCUU—GAA—U---GGD----UUC_GUGGGCUCAUPPCCCAUUA--------------CUAUAAAGTPCGAUUCUUUAAAGCGGCCCCA
================================================
```

63-72

### Tyr

```
>tdbR00000565|Bos_taurus|9913|Tyr|9PA
 CCUUCLAUALCUCAGDD-GGXU-AGAGCRRAGGACU9PAKAZCCUUAG--------------7D?GCUGGTPCG”UUCCGGCUCGAAGGACCA
>tdbR00000563|Lupinus_albus|3870|Tyr|GPA
 -CCGACCUUALCUCAGDD-#GU—AGAGCRGAGGACUGPAKAPCCUUAG--------------7XCACUGGUPCG”AUCCGGUAGGUCGGACCA
>tdbR00000562|Nicotiana_rustica|4093|Tyr|GPA
 -CCGACCUUALCUCAGDD-#GU—AGAGCRGAGGACUGPAKAPCCUUAG--------------7UCGCUGGUPCG”AUCCGGCAGGUCGGACCA
>tdbR00000557|Pichia_jadinii|4903|Tyr|GPA
 CUCUCGGUKLCCAAGDD-#GDDDAAGGCRPCAGACUGPA+APCUGAAC--------------AD?GGGCGTPCG”AUCGCCCCGAGAGACCA
>tdbR00000555|Saccharomyces_cerevisiae|4932|Tyr|GPA
 CUCUCGGUALCCAAGDD-#GDDDAAGGCRCAAGACUGPA+APCUUGAG--------------AD?GGGCGTPCG”CUCGCCCCGGGAGACCA
>tdbR00000554|Scenedesmus_obliquus|3088|Tyr|GPA
 -CCCCUUGUAGCUCAGDD-#GC—AGAGCRGGAGACUGPAKAPCUCUAG--------------7.CACUGGTPCG”AUCCGGUCGAGGGGACCA
>tdbR00000556|Schizosaccharomyces_pombe|4896|Tyr|GPA
 CUCCUGAUKGUGPAGDD-GGDD-AUCACACCCGGCUGPA+ACCGGUUG--------------7U?GCUAGTPCG”UUCUGGCUCAGGAGACCA
===================================================
```

### Asn

```
>tdbR00000304|Bos_taurus|9913|Asn|QUU
-GUCUCUGUKLCGCAADC-GGDX-AGCGCRPPCGGCUQUU6ACCGAAAG--------------7DUGGUGG.PCG”GCCCACCCAGGGACGCCA
>tdbR00000306|Homo_sapiens|9606|Asn|.UG
-GGUCCCAUKGUGPAAU--#GDD-AGCACUCPGGABU.UGAAPCCAGCG--------------A-U??GAGPPCA”AUCUCGGUGGGACCUCCA
>tdbR00000301|Lupinus_luteus|3873|Asn|;UU
 -GCCUCAGUALCUCAGD—GGDX-AGAGCRGUCGGBU;UU6ACCGAUAG--------------7D?GUAGGTPCG”GCCCUACUUGGGGCGCCA
>tdbR00000300|Saccharomyces_cerevisiae|4932|Asn|GUU
-GACUCCAUGLCCAAGDD-GGDD-AAGGCRUGCGACUGUU6APCGCAAG--------------AD?GUGAGTPCA”CCCUCACUGGGGUCGCCA
===================================================
```

### Cys

```
>tdbR00000621|Nicotiana_rustica|4093|Cys|GCA
 -GGGUCCAUAGCUCAGD--#GD—AGAGCAPPUGABUGCAKAPCAAGAG--------------7DCACCGGUPCG”ACCCGGUUGGGCCCUCCA
>tdbR00000021|Saccharomyces_cerevisiae|4932|Cys|GCA
  -GCUCGUAUGGCGCAGD—GGD—AGCGCAGCAGAPUGCA+APCUGUUG--------------7D?CUUAGTPCG”UCCUGAGUGCGAGCUCCA
===================================================
```

### Ile

```
>tdbR00000173|Bombyx_mori|7091|Ile|IAU
-GGCCCAUUALCUCAGDD-GGDN-AGAGCRPCGUGCUIAU6ACGCGAAG--------------7U?G?GGGTPCG”UCCCCUCAUGGGCCACCA
>tdbR00000172|Lupinus_luteus|3873|Ile|IAU
-GGCCCAUUALCCCAGDD-GGDD-AAGGCRPGGUGCUIAU6ACGCCGAA--------------GD?AGCAGTPCG”UCCUGCUAUGGGCCACCA
>tdbR00000174|Mus_musculus|10090|Ile|IAU
-GGCCGGUUALCUCAGDD-GGDD-AGAGCRPGGPGCUIAU6ACGCCAAG--------------7D?GCGGGTPCG”UCCCCGUACGGGCCACCA
>tdbR00000171|Pichia_jadinii|4903|Ile|IAU
-GGUCCCUUGLCCCAGDD-GGDD-AAGGCRPGGUGCUIAU6ACGCCAAG--------------AD?AGCAGTPCG”UCCUGCUAGGGACCACCA
>tdbR00000170|Saccharomyces_cerevisiae|4932|Ile|IAU
-GGUCUCUUKLCCCAGDD-GGDD-AAGGCACCGUGCUIAU6ACGCGGGG--------------AD?AGCGGTPCG”UCCCGCUAGAGACCACCA
===================================================
```

### Phe

```
>tdbR00000093|Bombyx_mori|7091|Phe|#AA
 -GCCGAAAUALCUCAGDD-GGG—AGAGCRPPAGABU#AAKAPCUAAAG--------------7D?CCUGGTPCA”UCCCGGGUUCGGCACCA
>tdbR00000089|Brassica_napus|3708|Phe|#AA
 -GCGGGGAUALCUCAGDD-GGG—AGAGCRPCAGABU#AAYAPCUGAAG--------------7DCGCGUGTPCG”UCCACGCUCACCGCACCA
>tdbR00000090|Lupinus_luteus|3873|Phe|#AA
 -GCGGGGAUALCUCAGDD-GGG—AGAGCRPCAGABU#AAWAPCUGAAG--------------7DCACGUGTPCG”UCCACGUUCACCGCACCA
>tdbR00000082|Neurospora_crassa|5141|Phe|#AA
 -GCGGGUUUALCUCHGDD-GGG—AGAGCRPCAGABU#AAYAP?UGAAG--------------7D?GUGUGTPCG”UCCACACAAACCGCACCA
>tdbR00000083|Saccharomyces_cerevisiae|4932|Phe|#AA
 -GCGGAUUUALCUCAGDD-GGG—AGAGCRCCAGABU#AAYAP?UGGAG--------------7UC?UGUGTPCG”UCCACAGAAUUCGCACCA
>tdbR00000084|Saccharomyces_cerevisiae|4932|Phe|#AA
 -GCGGACUUALCUCAGDD-GGG—AGAGCRCCAGABU#AAYAP?UGGAG--------------7UC?UGUGTPCG”UCCACAGAGUUCGCACCA
>tdbR00000081|Scenedesmus_obliquus|3088|Phe|GAA
 -GGCUUGAUALCUCAGCD-#GG—AGAGCRPPAGABUGAAKAPCUACAG--------------7.?CCCAGTPCG”U?BUGGGUCAGGCCACCA
>tdbR00000085|Schizosaccharomyces_pombe|4896|Phe|#AA
 -GUCGCAAU;LUGPAGDD-GGG—AGCAPLACAGABU#AAYAPCUGUUG--------------7NCAUCGGTPCGAUCCCGGUUUGUGACACCA
=================================================
```

### Thr

```
>tdbR00000443|Saccharomyces_cerevisiae|4932|Thr|IGU
 -GCUUCUAUGLCCAAGDD-GGD—AAGGCRCCACA’UIGU6APGUGGAG--------------AD?AUCGGTPCA”AUCCGAUUGGAAGCACCA
>tdbR00000444|Saccharomyces_cerevisiae|4932|Thr|IGU
 -GCUUCUAUGLCCAAGDD-GGD—AAGGCRCCACA’UIGU6APGUGGAG--------------AD?GUCGGTPCA”AUCCGACUGGAAGCACCA
===================================================
```

### Trp

```
>tdbR00000497|Bos_taurus|9913|Trp|BCA
 -GACCUCLUKLCGCAAD--#GD—AGCGCRPCUGABUBCAKAZCAGAAG--------------7DUGCGUGPPCG”AUCACGUCGGGGUCACCA
```
