## Supplementary material for "Coding triplets in the tRNA acceptor-TΨC arm and their role in present and past tRNA recognition": Table S1

**Table S1a- Occurrence statistics- random, sample and individual distributions of bacterial data of conserving amino acids†**

| Amino acid (no. sequences) | Obs.* | Random nucleotide distribution (A,C,G,T)= (25,25,25,25)% | | | Individual nucleotide distribution | | | |
|---|---|---|---|---|---|---|---|---|
| | | **Exp.** ** **(%)** ±0.8 | **Exp.** ** **(No.)** | ***P****** | **Distribution (A,C,G,T) %** | **Exp.** ** **(%)** ±0.8 | **Exp.** ** **(No.)** | ***P****** |
| **Conserving amino acids : number of observed coding triplets >> number of expected coding triplets** | | | | | | | | |
| Ala (209) | 174 | 24% | 50 | $10^{-89}$ | (8,46,19,27) | 45% | 94 | $10^{-28}$ |
| Asp (110) | 106 | 12% | 13 | $10^{-166}$ | (7,48,25,20) | 18% | 20 | $10^{-100}$ |
| Gly (243) | 229 | 19% | 46 | $10^{-197}$ | (5,54,19,22) | 74% | 180 | $10^{-12}$ |
| His (83) | 75 | 22% | 18 | $10^{-51}$ | (22,44,14,20) | 41% | 34 | $10^{-20}$ |
| Pro (180) | 177 | 12% | 22 | $10^{-272}$ | (7,49,20,24) | 35% | 63 | $10^{-70}$ |
| Ser (290) | 286 | 33% | 96 | $10^{-124}$ | (5,62,3,30) | 87% | 252 | $10^{-9}$ |
| Arg (305) | 291 | 59% | 181 | $10^{-37}$ | (9,32,39 20) | 83% | 254 | $10^{-8}$ |
| Leu (366) | 307 | 51% | 187 | $10^{-36}$ | (4,49,28,19) | 61% | 222 | $10^{-19}$ |
| Val (201) | 176 | 71% | 143 | $10^{-7}$ | (16,43,20,21) | 65% | 131 | $10^{-11}$ |

† Statistics for the sample distribution of the conserving is given in Tab.2 of the manuscript.

* Obs. - For the conserving amino acids- number of pre-3'end strings holding at least one conserved cognate C/AC.

** Exp. - percentage and number of pre-3'end sequences expected to carry a conserved cognate C/AC. Expectation values were derived according to the algorithm described in Materials and Methods.

*** ***P*** - The probability that the observed occurrence of a particular C/AC set is random.

**Table S1b- Occurrence statistics- random, sample and individual distributions of bacterial data of non-conserving amino acids**

| **Amino acid (no. sequences)** | **Obs.** | **Random Distribution** (A,C,G,T)= (25,25,25,25)% | | | **Individual distributions** | | | | **Sample Distribution** (A,C,G,T)= (10,46,24,20)% | | |
|---|---|---|---|---|---|---|---|---|---|---|---|
| | | **Exp. (%)** ±0.8 | **Exp. (no.)** | ***P*** | **Distribution (A,C,G,T) %** | **Exp. (%)** ±0.8 | **Exp. (no.)** | ***P*** | **Exp. (%)** ±0.8 | **Exp. (no.)** | ***P*** |
| **Amino acids with number of observed coding triplets << number of expected coding triplets** | | | | | | | | | | | |
| Gln (120) | 10 | 44% | 53 | $10^{-15}$ | (22,45,17,16) | 40% | 48 | $10^{-12}$ | 35% | 42 | $10^{-9}$ |
| Glu (113) | 25 | 42% | 47 | $10^{-5}$ | (12,28,42,18) | 35% | 40 | $10^{-3}$ | 45% | 51 | $10^{-6}$ |
| Lys (122) | 0 | 35% | 43 | $10^{-16}$ | (16,44,23,17) | 18% | 22 | $10^{-7}$ | 19% | 23 | $10^{-7}$ |
| Met (218) | 8 | 21% | 46 | $10^{-10}$ | (12,46,27,15) | 9% | 20 | $10^{-3}$ | 9% | 20 | $10^{-3}$ |
| Thr (169) | 67 | 71% | 120 | $10^{-19}$ | (12,32,35,21) | 66% | 112 | $10^{-13}$ | 58% | 98 | $10^{-6}$ |
| Tyr (98) | 2 | 33% | 32 | $10^{-10}$ | (8,66,5,21) | 11% | 11 | $10^{-3}$ | 12% | 12 | $10^{-3}$ |
| **Amino acids with number of observed coding triplets ≈ number of expected coding triplets** | | | | | | | | | | | |
| Asn (108) | 10 | 42% | 45 | $10^{-12}$ | (23,21,41,15) | 24% | 26 | $10^{-4}$ | 15% | 16 | **NS** |
| Cys (80) | 16 | 40% | 32 | $10^{-4}$ | (9,43,39,9) | 26% | 21 | **NS** | 32% | 26 | **NS** |
| Ile (133) | 16 | 42% | 56 | $10^{-12}$ | (19,49,18,14) | 18% | 24 | **NS** | 14% | 19 | **NS** |
| Phe (90) | 4 | 35% | 32 | $10^{-10}$ | (7,43,36,14) | 9% | 8 | **NS** | 19% | 17 | $10^{-4}$ |
| Trp (86) | 21 | 23% | 20 | **NS** | (4,54,14,28) | 13% | 11 | $10^{-3}$ | 25% | 22 | **NS** |

* Obs. - number of sequences holding at least one cognate codon or anticodon.
** Exp. - percentage and number of sequences expected to carry any cognate C/AC. Expectation values were derived according to the algorithm described in Materials and Methods.
*** ***P*** - The probability that the observed occurrence of a particular C/AC set is random.
**NS** - Statistically non-significant ($P \geq 0.01$)
