## Supplementary material for "Coding triplets in the tRNA acceptor-TΨC arm and their role in present and past tRNA recognition": Table S2

**Table S2 – conserved C/ACs occurrence in the pre-3'end of different bacterial genera.**

Data for the robustly conserving amino acids from the most populated bacterial genera, i.e. *Bacillus* (278 gene-sequences, gram-positive, phylum Firmicutes, 6 species) and *Escherichia*, (252 gene-sequences, gram-negative, phylum Proteobacteria, 1 specie), Salmonella (148 gene-sequences, gram-negative, phylum Proteobacteria, 2 species) and *Mycoplasma* (231 gene-sequences, phylum Tenericutes, neutral in staining, 6 species). While the first three bacteria exhibit complete conservation of the C/ACs in the pre-3'end sequences, the occurrence significantly deviates in *Mycoplasma*.

| Conserving amino acids | *Bacillus* [(no. strings) | *Escherichia* (no. strings) | *Salmonella* [(no. strings) | *Mycoplasma* [(no. strings) |
|---|---|---|---|---|
| **Ala** | 100 % (10) | 100 % (11) | 100 % (6) | 0 % ( 8) |
| **Asp** | 100 % (11) | 100 % ( 6) | 100 % (3) | 43 % ( 7) |
| **Gly** | 100 % (22) | 100 % (15) | 100 % (9) | 8 % (13) |
| **His** | 100 % (12) | 100 % ( 5) | 100 % (4) | 67 % ( 6) |
| **Pro** | 100 % ( 9) | 100 % (15) | 100 % (10) | 88 % ( 8) |
| **Ser** | 100 % (25) | 100 % (20) | 100 % (13) | 100 % (22) |
